# Kleptocytosis: Comparative Proteomics and Functional Imaging Show *Varroa destructor* Co-opts Intact Host Proteins for Rapid Development and Reproduction

**DOI:** 10.1101/2022.09.30.509900

**Authors:** Lincoln N. Taylor, Justin D. Hein, Daniel Sonenshine, Reid Panzer, Mikayla Henry, Chris Borke, Jackson Kominsky, Wayne O. Hemphill, Steven C. Cook, Madison Sankovitz, Connor Gulbronson, Jay Evans, Dennis van Engelsdorp, Chris Ebmeier, Ron Ochoa, Samuel D. Ramsey

## Abstract

*Varroa destructor* must produce mature offspring within the inflexible nine-day window framing the pupal development of their honey bee host. Missing this deadline renders the foundress mite’s fitness zero, establishing evolutionary pressure to accelerate reproduction and development. Through micro-computed tomography and modeling *Varroa’s* energy budget, we found each egg grows to constitute ∼18% of the foundress’s body volume in 30 hours, yet accounts for less than 10% of her energy usage. We hypothesized that this small energy investment is a successful strategy because of *Varroa*’s long-ignored ability to traffic intact host proteins. Through gel electrophoresis, concomitant immunodetection, and MS/MS, we detected several intact, bee-derived proteins in mite eggs, including metamorphic proteins (hexamerins) and egg-yolk precursors (vitellogenin and apolipophorin) which likely reduce the parasite’s direct reproductive investment. We then expressed recombinant Halotag labelled *Apis* vitellogenin to determine the route conveying exogenous proteins into the mite’s oocyte. We detected fluorescent vitellogenin in the lyrate organ and observed a tube-like extension of the lyrate organ connecting to the ovum, likely providing an avenue for intact host proteins. Finally, we tested the hypothesis that exogenous proteins maintain functionality in the parasite. *Varroa* metamorphose despite their inability to produce hexamerins, critical metamorphic proteins. Through label-free quantification of proteins in metamorphosing mites, we observed a hexamerin depletion pattern consistent with usage as a metamorphic amino acid reservoir. We describe this process as “kleptocytosis,” denoting movement of stolen intact macromolecules from host to a parasite cell. Given their fixed developmental timeline, this pathway presents a promising target for novel *Varroa* management strategies.

**Significance Statement:** Global honey bee health is threatened by the parasitic mite Varroa destructor; its success due in part to its rapid reproduction and accelerated development. By combining biological modeling, micro-computed tomography, fluorescence imaging, and quantitative proteomics, we show that Varroa siphon intact, ostensibly functional host proteins conveying them directly to their oocyte. This avoids the energetically inefficient process of digesting and reconstructing ingested proteins and affords the capacity to utilize proteins that it does not produce de novo facilitating rapid reproduction and accelerated development. We call this process “kleptocytosis,” and identify anatomical adaptations which apparently facilitate protein movement. This work exposes a new target in Varroa’s physiology, providing a promising direction for more effective management strategies.

## 1. Introduction

Mites (Class Arachnida; Subclass: Acari) have evolved to exploit nearly every terrestrial ecological niche, owing primarily to their diminutive size, efficient resource processing, and inventive reproductive strategies (1). The parasitic mite *Varroa destructor* (Order: Mesostigmata), hereafter “*Varroa,*” has adapted to exploit the western honey bee, *Apis mellifera* (Order: Hymenoptera), to a degree of success rarely seen in other parasites (2, 3). In less than 100 years since its transition from its original host, *Apis cerana*, this parasite has achieved a cosmopolitan distribution and is a ubiquitous pest of beekeeping operations (4, 5). Global economic dependence on commercially pollinated crops has led to frequent and expensive interventions to manage the deleterious impact of this parasite on honey bee health (6–8). The increasing virulence of this parasite, coupled with its ubiquity in beekeeping operations, underscores the imperative to understand the factors contributing to its success (9, 10). Chief amongst these is *Varroa*’s ability to synchronize its reproductive and developmental rates with the strict developmental time schedule of its metamorphosing host.

The developmental timeline of the larval bee host strictly constrains *Varroa* fitness (2). For reasons undetermined in previous work, *Varroa* can only develop through their immature stages and reproduce while feeding on immature bees (11). The host’s rapid 12-day transition from prepupa to adult compounded by the extended gestation required for the initial male *Varroa* egg (70 hours) constrains the foundress to a narrow nine-day window for producing mature, mated offspring (12). Mites that fail to fully develop and mate during this short period have a fitness of zero (2, 13), demonstrating an evolutionary pressure for rapid reproduction and development.

Prior to reproduction, *Varroa* enter a stage wherein they feed repeatedly on adult bees (primarily nurse bees) for an average of seven days or as long as two weeks (14). Mites prevented from feeding on adult bees present high rates of infertility, suggesting that nutrients critical for reproduction are acquired during this time (14). After invading a brood cell, the *Varroa* foundress produces a relatively large egg every 30 hours (15) (**Fig. 1*A-D***). Additionally, development within the egg is accelerated. While other Mesostigmata hatch as a six-legged larva and molt into an eight-legged protonymph outside the egg, this process occurs rapidly inside the ovum in *Varroa*, functionally skipping the larval stage and instead hatching as an eight-legged protonymph (**Fig. 1*C***) (16). Another life stage common in Mesostigmata, the tritonymph, is also bypassed in favor of rapid metamorphosis from deutonymph directly to the adult stage (16).

Prior to each molt, and ostensibly within the egg, *Varroa* enter an immobile stage, unresponsive to stimuli (12, 15). These stages have been described as calyptostase–periods of rapid metamorphosis within the chorion or the cuticle of the previous instar– on the basis of observed quiescence alone. This is without proof of rapid physiological reorganization (2, 15). While calyptostase has been observed in other Acari, it has not been formally documented in other Mesostigmata, aside from the family Varroaidae (17, 18). In other parasitic mites, it is hypothesized to have evolved as a means to synchronize developmental timelines with their hosts and to facilitate dramatic morphological changes between instars (e.g. the development of new appendages or reproductive organs) (1, 18). A calyptostase is analogous to an insect pupa (18). As such, calyptostatic mites are subject to the same biochemical challenges of rapidly metamorphosing insect pupae, which require storage proteins with all essential amino acids necessary to fuel production of the new tissues comprising the imago. Insects meet this demand by producing complex storage proteins called hexamerins, which are broken down to provide the expansive pool of amino acids needed to complete metamorphosis (19). *Varroa* do not produce these proteins, thus their capacity for rapid metamorphosis demands explanation (20). Rapid egg production likewise requires a substantial biochemical investment, primarily through vitellogenins (Vgs). Vitellogenins are large lipid transfer proteins which act as precursors to vitellin, the primary egg-yolk protein (21), and serve to provision the developing embryo. The number of different Vgs produced is species-dependent; honey bees only produce one vitellogenin, while *Varroa* produce two Vgs and a Vg-like protein. The metabolic cost of *de novo* synthesis for these high-molecular-weight proteins (210-213 kDa) creates a significant resource hurdle (22, 23).

While mites have access to an abundant and nutritionally dense food source in the partially digested tissue aggregate in developing bees’ hemocoel (24, 25), prior research by Tewarson & Engels (1982) suggests that *Varroa* are inefficient at processing these nutrients, evidenced by numerous undigested and intact *Apis*-host proteins found in mite feces (26). Thus, further explanation is warranted as to how these mites can sustain their observed rapid developmental and reproductive rates. Static energy budgets have been used previously to model energy allocation based on the conservation of energy, where energy inputs are equal to the energy outputs (27), and thus are a useful tool for determining how much energy *Varroa* are allocating to reproduction versus other processes.

In addition to the intact proteins found in the feces, Tewarson & Engels (1982) also observed unidentified proteins in *Varroa* that migrated at the same rate as host proteins via gel electrophoresis, suggesting that *Varroa* could pass host proteins through their bodies intact (26). This phenomenon is a rare strategy but not without precedence; other medically significant parasites, including tsetse flies and ticks, transport large and intact exogenous proteins across different tissues (28, 29). However, provisioning an ovum with intact host proteins would be without precedence. Tewarson and Engels (1982) proposed that *Varroa* move host proteins into their eggs, though without determining the identity of these proteins, without confirming they were of host origin, or providing a mechanism for how they could be trafficked into the egg (26,30). Because egg components are typically acquired through highly specific receptor-mediated transport (22, 31), the utilization of mite-specific pathways by host *Apis* proteins is highly improbable. Without a mechanism for foreign proteins to enter the egg from the digestive system, subsequent detections of *Apis* proteins in *Varroa* eggs were previously dismissed as contamination (20), and the ability of *Varroa* to provision their eggs with co-opted host proteins has not been researched since.

We hypothesized that the host proteins detected in *Varroa* eggs are not representative of contamination but are crucial for the mite’s growth and accelerated development within the egg. We predicted that the detected proteins would be those involved in nutrient mobilization. These foreign proteins would need to be introduced through means that bypass normal receptor-mediated transport. We further expected that stolen host proteins sequestered in immature mites would support the metamorphic processes underpinning the capacity of *Varroa* to rapidly metamorphose between life stages. To address these hypotheses: (**1**) we constructed the static energy budget for reproductive *Varroa* using bomb calorimetry to determine whether foundress mites enter a period of higher metabolic efficiency which can account for their capacity to rapidly produce large, advanced ova. (**2**) Via mass spectrometry, we worked to determine if foreign proteins are utilized by reproductive *Varroa* mites, whether they are taken up selectively by the ova, and the identities of these co-opted host proteins (**3**) By engineering a recombinant HaloTag-labeled host protein, we used confocal microscopy to investigate how *Varroa* could traffic host proteins to the oocyte while bypassing conventional receptor-mediated systems; these observations were complemented by micro-computed tomography (microCT) to visualize the associated internal reproductive anatomy. (**4**) Finally, we tested the hypothesis that *Varroa* utilize host hexamerins to fuel metamorphosis. We analyzed hexamerin titers during, before, and after calyptostase via label-free quantification to determine whether the depletion pattern was consistent with usage as a metamorphic reservoir.

## 2. Materials and Methods

### Sample collection

Mites, mite eggs, and bees were collected from apiaries at the USDA-ARS, Beltsville, MD, Boulder, CO, and Delray Beach, FL. Dispersal-phase *Varroa* were collected from adult bees with a paintbrush or powdered sugar. Specimens collected with powdered sugar were washed to remove traces of sugar and cornstarch. To access reproductive and immature *Varroa*, we opened capped cells with straight dissecting forceps. Once opened, reproductive mites and their eggs were collected with a paintbrush, avoiding contact with the host bee. Immature mites, including deutonymphs, calyptostase, and teneral (*i.e.*, newly molted adult) mites were also collected (see Fig. S6 for examples of these mite stages). We used separate paintbrushes for each stage and cleaned paintbrushes with 70% ethanol between specimen collections. We also collected *Varroa* feces from opened brood cells, using a metal pin to pierce the solid feces pile, then angling the pin to dislodge the feces from the wall of the cell.

To collect the fat body-hemolymph aggregate from immature bees, we opened capped brood cells and pushed wax away from the sides to expand the hole as much as possible to ensure the opening could accommodate the bee’s removal without rupture. If the pupa was ruptured during this process, it was discarded and nothing else from the cell was collected. We extracted the fat body–hemolymph aggregate from pupae via a positive displacement pipette, following a small incision down the side of the pupal abdomen with ophthalmic surgery micro scissors (Moria Surgical). Gut tissue was removed first with the positive displacement pipette under standard dissecting scope magnification before the dissociated fat body aggregate was collected in the fluid matrix of the hemolymph. Parietal fat body collected from the metasoma of dissected adult bees was removed with fine-tipped forceps. Dissections were performed under standard dissecting scope magnification with phosphate buffered saline (PBS). We collected *Apis* eggs from uncapped cells using a metal grafting tool and *Varroa* eggs using a fine depilated paintbrush. All samples were immediately stored at −80°C until further processing.

#### 2.1.1 Static energy budget

We created a static energy budget for reproductive *Varroa destructor* outlining the 30-hour period it takes to gestate a single egg. We used the following equation for our energy budget modified for *Varroa* from the standard equation (27):

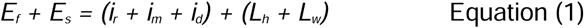

Where the energy income into the system is represented by *E_f_* (energy from ingested food) and *E_s_* (energy stored while feeding). Representing the entirety of the energy pool the mites have to draw from, it must be equal to their investments in reproduction (*i_r_*) & maintenance (*i_m_*) (e.g. energetic costs associated with basal metabolic rate, cellular upkeep, homeostasis & internal regulation, immune function, structural repair, etc.). In addition, energy is lost via respiration in the form of heat (*L_h_*) and waste (*L_w_*). Dynamic energy budgets include a term to account for energy inputs to growth. However, as we are focusing on the reproductive phase in *Varroa* foundresses, we can omit this as energy dedicated to growth and development is zero in adults. We omit this term (*i_d_*) and assume that all available energy is used solely for reproduction and maintenance (32) or put simply:

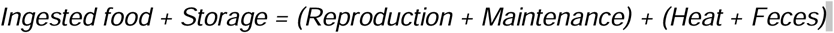

We assume that storage is encompassed by the energy a mite accumulates during the dispersal stage, or the time between emerging from and re-entering a new cell for reproduction. As such, we collected mites that had just entered a brood cell but had not yet fed; this was determined by selecting mites located in the brood food, which only occurs prior to the host’s cocoon spinning stage, when *Varroa* first feed (33). We also monitored cells with emerging bees to collect newly emerged mites before they could feed on an adult host. The difference between the energy in these two groups of mites, representing the beginning and end of the dispersal phase (assumed to be at full storage and storage depletion), allows us to determine how much energy is stored during the dispersal phase. We cannot fully account for the potential that *Varroa* store some level of energy while feeding on their reproductive host. However, given the energetically taxing nature of reproduction, we assume that this is a period of intense energy output, and energy inputs into storage during this stage are negligible. As such, the storage value represents the energy acquired solely during the dispersal stage. Finally, this model assumes the constant feeding rate (1 µL/day) and deposition rate of 106.2 µg feces/day (34). To measure the energy lost as heat, we placed 177 gravid mites into a sealed inert glass container with a high precision temperature sensor (TMP117 ± 0.1°C, Texas Instruments), which was queried each second (***SI Appendix* S1**).

We used bomb calorimetry to analyze the energy content of the inputs and outputs of our equation. We analyzed gravid mites (253 mites, 0.100 g total), *Varroa* eggs (300 eggs, 0.01 g total), pupal *Apis* fat body–hemolymph aggregate (0.2 g), and *Varroa* feces (0.06 g). Additionally, we analyzed mites from cells that had recently been capped, prior to their reproductive stage (96 mites, 0.047 g total), and mites that were emerging from brood cells (100 mites, 0.036 g total). Samples were washed in ethanol to ensure that no debris or brood food components inflated energy values. All samples were stored at −80°C prior to shipping for bomb calorimetry, which was performed at the Weill Cornell Medicine (WCM) Mouse Phenotyping Core (Medical College of Cornell University, NY, USA) (***SI Appendix* S2**).

#### 2.1.2 Estimating the translational capacity of a *Varroa* foundress

We estimated the translational capabilities of *Varroa* for Vg synthesis to assess the challenge that unassisted egg provisioning would present. We based our calculations on *Varroa* Vg2, a 210 kDa polypeptide consisting of 1,835 amino acid residues (23). We then estimated the amount of *Varroa* Vg to be 90% of the protein content in an egg, assuming similar protein composition to the eggs of other Parasitiformes (e.g., ticks) (35–37). This yields an estimated 3.5 x 10^13^ Vg molecules per egg. To approximate how many ribosomes would be needed to support the required amount of Vg production, we estimated the total number of available ribosomes within a foundress *Varroa*. Where *N_Vg_* is the total number of synthesized Vg molecules, *N_r_* is the number of available ribosomes, *T* is the total gestation time (30 hours), *t*_elongation_ is the time needed to produce one Vg molecule.

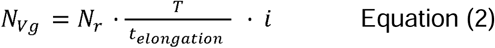

Single-molecule imaging studies have demonstrated that translation occurs in bursts rather than as a constitutive process (38–40). Because the initial calculation assumes that Vg transcripts are continuously available to the foundress mite’s translational machinery throughout the 30-hour reproductive period, we will treat the above calculation as an unlikely biological ceiling in this process. To establish a realistic comparison, we then introduced a translational inefficiency factor correction factor (*i*) to account for intermittent translation. No data currently exist describing *Varroa* Vg transcript abundance or burst kinetics in foundress mites, but we know that *i* < 1. As such, we have selected *i* =0.8 as a conservative value, representing a system in which transcripts are productively engaged in translation for approximately 80% of the available reproductive interval, biasing the model towards overestimating rather than underestimating the endogenous translation capacity of the mite.

Eukaryotic elongation rates can vary from 3-5 amino acids per second, depending on the mRNA transcript (41, 42). We first estimate the number of ribosomes needed to synthesize the required amount of Vg assuming an optimal translation rate of 5 aa s^-1^ (*t_elongation_* = 367) and idealized translational efficiency (*i* = 1). We then estimate the ribosomal requirement under more realistic parameters. We further evaluate less than optimal elongation rates of 4 aa s^-1^ (*t_elongation_* = 459), and 3 aa s^-1^ (*t_elongation_* = 612), as well as translational discontinuity (*i* = 0.8).

#### 2.2.1 Protein Resolution via Gel Electrophoresis and Immunodetection of *Apis-*Vg

*Varroa m*ites (n = 20-30 reproductive, 20-30 dispersal), *Varroa* eggs (n = 15-20), *Apis* eggs (n = 10), and tissue samples collected from *Apis* pupae (n = 5) and adult bees (n = 5) were washed in ultra-pure water (Invitrogen) and placed in separate 1 ml microcentrifuge tubes with 150 µL of the appropriate buffer, a non-ionic lysis buffer (150 mM NaCl, 0.1% Triton-X 100, 50 mM Tris-HCl) or NativePAGE Sample Buffer (Bio-Rad). Samples were homogenized in the lysis buffer with 0.5mm glass disruption beads (Thermo Fisher Scientific) using a benchtop tissue homogenizer (Fast Prep-24, MP Biomedicals, Santa Ana, CA) for three 30-second intervals. Samples were then centrifuged (4°C) at 15,000 x g for 10 minutes. Afterward, the total protein concentration of the samples was assessed using a Nanodrop model 8000 (Thermo Fisher Scientific).

Two SDS-PAGE gels (Invitrogen NuPage 4-12% Bis-Tris Mini Protein Gels) were run simultaneously with 50 µg per lane at 200V (50 minutes). One gel was stained with Coomassie R-250, and the other gel was left unstained for western blot immunodetection of *Apis* vitellogenin. Proteins were transferred onto a 0.45 µm pore PVDF membrane (Invitrogen). The membrane was incubated in blocking solution, rinsed, and incubated in anti-*A. mellifera* Vg (produced by Custom Antibodies Thermo Fisher Scientific) diluted to 1:2500 for 1 hour. Blotting was conducted with the anti-rabbit WesternBreeze chromogenic detection system (Invitrogen). Details on antibody generation can be found in ***SI Appendix* S3**. For comparison, NativePAGE gels were run to determine whether the proteins observed were structurally intact.

#### 2.2.2 Mass Spectrometry on Candidate Bands

Candidate bands for proteins sequestered in mite eggs were evaluated by comparing *Apis* tissue lanes and *Varroa* egg lanes on the Western blot. The candidate bands excised from the SDS-PAGE gel included adult *Apis* fat body, pupal *Apis* fat body–hemolymph aggregate, dispersal-phase *Varroa*, reproductive-phase *Varroa,* and V*arroa* eggs. The gel slices were placed in separate microcentrifuge tubes with a wet paper towel to prevent dehydration and shipped to the Keck Biomolecular Research Facility at the University of Virginia (Charlottesville, VA) for analysis by MS and MS/MS ***(SI Appendix* S4)**. Each gel band was run as a unique sample. The resulting data were analyzed using database searching with the Sequest search algorithm (Thermo Fisher Scientific; version IseNode in Proteome Discoverer 2.2.0.388) against NCBI *Apis mellifera* and *Varroa destructor* sequences. All peptides and proteins were thresholded at a 1% false discovery rate (FDR). Spectral counts were normalized in Scaffold (Proteome Software) using the standard spectrum count normalization algorithm, in which spectral counts for each sample were scaled by the ratio of the mean total spectral count across all samples to the total spectral count of the individual sample. Total spectral counts for each protein within a given tissue type were calculated by summing normalized spectral counts from each constitutive gel band sample.

#### 2.3.1 Micro-Computed Tomography (MicroCT) Scans of *Varroa*

To identify possible pathways for foreign proteins to enter the mite’s ovum, we scanned gravid female *Varroa* using microCT imaging. Samples were stained with iodine-iodide. Mites were scanned at the Mouse Imaging Facility at the National Institute of Neurological Disorders and Stroke (Bethesda, MD), using a Bruker SkyScan 1172 microCT scanner. MicroCT imaging was also used to determine the volume for adult *Varroa* and their eggs **(*SI Appendix* S5)**.

#### 2.3.2 Vg-HaloTag-Fluorophore Synthesis and *in vivo* feeding trials

After consistently finding intact or nearly intact *Apis* Vg in mite eggs, we selected this protein as a candidate protein to track the mite’s post-ingestion pathway through fluorescence microscopy. We engineered a Vg-HaloTag construct, which is compatible with multiple fluorophores. Total RNA was extracted from *A. mellifera* worker fat body (Direct-zol RNA MiniPrep Kit, Zymo Research) and reverse-transcribed into DNA using random hexamer primers (Superscript IV Reverse Transcriptase, Invitrogen). We amplified three overlapping 1.8 kb fragments of the vitellogenin coding sequence (NCBI RefSeq NM_001011578.1) (***SI Appendix* Table S1**). Isolated vitellogenin DNA fragments were cloned into pCR4-TOPO TA sequencing vectors (TOPO TA Cloning Kit, Thermofisher) and transformed into competent 5-alpha *E. coli* cells (New England Biolabs). The fragments were then linked into a new plasmid construct via Gibson assembly. This construct included a hexahistidine and maltose-binding protein (MBP) fusion with a Prescission protease cleavage site on the N-terminal end of the Vg coding sequence, as well as a Halo protein fusion on the C-terminal end. The plasmid was used to make an infectious baculovirus stock in Sf9 (*Spodoptera frugiperda*, IPLB-Sf-21-AE) cells using the Bac-to-bac system (Invitrogen). To express recombinant protein, High Five (*Trichoplusia ni*, BTI-Tn-5B1- 4) cells were transfected with baculovirus at 28°C for 66 hours, frozen in liquid nitrogen, and stored at -80°C for Vg purification (***SI Appendix* S6**). Following purification, incubation with the JF549 fluorophore (Janelia Fluor Dyes) allowed covalent binding to the HaloTag (***SI Appendix* S7**).

The Vg-HaloTag-JF549 construct was introduced into live foundress *Varroa* mites in 48-hour feeding trials (mites collected from uncapped brood cells). We 3D printed a capsule to mimic a bee larva. One side of the capsule was open and filled with 100 µL of homogenized larval fat body and sealed with a parafilm membrane (***SI Appendix***, **Fig. S1*A***). The capsule was sealed using parafilm stretched to 15 µm thickness to mimic a bee’s membrane. To encourage feeding, we rubbed a bee pupa on the parafilm membrane (25). Mites in the control group were given only the homogenized fat body without fluorophore. For mites in the experimental group, we added 5 µL of Vg-HaloTag-JF549 (500 µM) as a layer on top of the fat body rather than homogenizing the mixture to increase the likelihood of ingestion. The capsule was then moved inside a 2 mL tube and attached to the inner wall using beeswax (***SI Appendix,* Fig. S1*B***).

We moved one to three mites into each tube and sealed it using plastic wrap for ease of observation. In total, we used nine mites in the control group and eight mites in the experimental HaloTag-fed group. Mites were kept in an incubator (34°C, 70% RH) throughout the trial. At 24 hours, the old food capsule was replaced, and the experimental group received another 5 µL HaloTag-Vg with the diet. After 48 hours, all mites were removed from their capsules, flash frozen in liquid nitrogen, and stored at -80°C. Prior to imaging, we removed the genital plate to improve visualization of the internal structures **(*SI Appendix*, Fig. S2)**. Fluorescence microscopy was performed at the BioFrontiers Institute’s Advanced Light Microscopy Core using a Nikon AXR Laser Scanning Confocal microscope. Mites were placed in Ibidi Imaging Dishes (Ibidi GmbH), ventral side down. 4X and 10X objectives were used depending on the number of mites. Imaging parameters (dwell time and averaging) were optimized to minimize photobleaching while maintaining sufficient signal-to-noise ratios to investigate fluorophore localization. Three laser lines were used in the scans: EGFP, TRITC, and AF647, to ensure the broadest possible detection of autofluorescence and fluorophore. Fluorophore-specific signals from the images were distinguished from similar autofluorescent spectra emanating from the mite’s exoskeleton via decomposing the images into signal matrices in R and processing the matrices through an algorithm to identify per-pixel correlations to fluorophore-like signal (***SI Appendix,* S8)**.

#### 2.4.1 Measuring Hexamerin Titers Across *Varroa* Development

We tested the hypothesis that *Apis* hexamerin, another protein family detected by MS in *Varroa* eggs, would be rapidly metabolized by *Varroa* during the calyptostatic stage. We measured hexamerin titers across three developmental stages: deutonymphs, calyptostatic deutonymphs, and teneral (newly molted) adults (see Fig. S6 for a representative image of each stage). We selected these stages such that hexamerins could be measured before, during, and after calyptostase. We prepared four samples of each of the three developmental periods, each sample consisting of two mites in a 1.5 ml tube. Prior to homogenization, the mites were washed in 200 µL sterile phosphate-buffered saline (PBS) by inverting the tube and pipetting out the liquid. We homogenized the samples in 50 µL of SDS buffer at room temperature using a sterile disposable pestle against the tube, then washed any residue off the pestle into the tube using another 50 µL SDS buffer. All samples were then incubated at 70°C for 10 minutes to denature the peptides. Equal homogenization for each sample was verified by silver stain. Samples were analyzed at the Proteomics and Mass Spectrometry Core Facility (RRID: SCR_ 018992) in the Department of Biochemistry at the University of Colorado Boulder for label-free quantification (LFQ) (***SI Appendix* S9**). Raw files were searched against the Uniprot *Varroa destructor* (UP000594260) and *Apis mellifera* (UP000005203) databases using Maxquant (version 2.6.3.0) with cysteine carbamidomethylation as a fixed modification. Methionine oxidation and protein N-terminal acetylation were searched as variable modifications. All peptides and proteins were thresholded at a 1% false discovery rate.

We identified 4,546 peptides across the three tested life stages (4,027 from *Varroa*, 480 from *Apis*, 39 peptides shared high similarity between both species). Log_2_ fold-changes in LFQ intensities for each protein, relative to their abundance in the previous life stage, were compared between deutonymphs, calyptostases, and teneral adults with Limma. A Benjamini-Hochberg adjustment was applied to all P-values (43). To determine whether hexamerins decreased to a greater extent than the background proteins during calyptostase, we filtered our dataset to include only *Apis* proteins that were observed across all three *Varroa* life stages. This resulted in a data subset containing 209 *Apis* proteins. We then separated hexamerin 110, 70b, and 70c from the other proteins (*n* = 206). Using a one-tailed permutation test with 100,000 permutations, we tested the hypothesis that these hexamerins would exhibit a mean decrease (measured by mean LFC) greater than that of other proteins. We compared the mean log_2_ fold-changes of these hexamerins to that of the other proteins for all three comparisons (deutonymphs to teneral, deutonymphs to calyptostase, and calyptostase to teneral).

## 3. Results

### 3.1 Static energy budget shows temporal and energetic constraints on *Varroa* foundresses

We determined the energy inputs and outputs during the 30-hour reproductive window through bomb calorimetry (***SI Appendix,* Table S2**), using Equation (1).

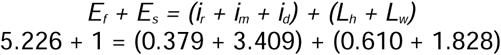

Over this 30-hour period, *Varroa* ingest 5.226 J of host fat body–hemolymph aggregate. In combination with the 1 J accumulated during the dispersal phase, this results in a total of 6.226 J of energy inputs. Over the 30-hour period, *Varroa* lose 1.828 J to feces, as well as an additional 0.61 J to heat. Our data were similar to *Varroa* heat-loss data from Garedew *et al.* (2004), which we extrapolated from their dataset to 0.54 J over 30 hours (44). Of the remaining 3.788 J, 9.99% is allocated to egg production (0.379 J). By solving for the remaining value of organismal maintenance, we estimate that *Varroa* dedicate about 3.409 J to bodily processes required to keep the organism alive (**Fig. 1*E***). Given that *Varroa* mites only gestate one egg per 30-hour period, this represents the mite’s entire reproductive energy investment per reproductive event. As such, our equation shows an energy budget shifted towards maintenance and waste, with the lowest investment in reproduction out of all categories. Despite this, our volumetric microCT scans revealed that, on average, an egg represents approximately 18% of the foundress’s body volume (***SI Appendix,* Table S3**).

We then estimated that the number of ribosomes needed for *Varroa* to produce the necessary amount of *Varroa* Vg2 in an egg during the 30-hour gestation period is likely beyond the translational capability of a *Varroa* foundress. We used *Varroa* Vg2, the most abundantly detected vitellogenin in mite eggs (**Table 1**). Using Equation (2), we first assumed an optimal elongation rate of 5 aa s^-1^ and an *i* = 1. Under these conditions, *Varroa* would require 1.2 *x* 10^11^ constantly active ribosomes. We then estimated the required number of ribosomes under more realistic scenarios. Accounting for discontinuous translation (*i* = 0.8), we then estimated the number of ribosomes needed for the range of elongation rates 5, 4, and 3 aa/s. This yielded 1.5 *x* 10^11^, 1.9 *x* 10^11^, and 2.5 *x* 10^11^ ribosomes, respectively.

To determine whether a foundress can meet that translational need, we estimated the total number of ribosomes available to the mite. Ribosome abundance is highly conserved in eukaryotic cells (10^5^ to 10^7^ ribosomes per cell) (45); using the upper limit, we estimate the number of available ribosomes in a foundress *Varroa* by scaling to body mass (0.49 mg), yielding approximately 10^12^ total ribosomes (46). We assume 75% of ribosomes are estimated to be actively engaged in translation at a given time (45, 47–50). Thus, we estimate that a foundress has approximately 9 x 10^11^ ribosomes available for producing other peptides. Under perfect conditions, a *Varroa* foundress would need to use 13% of its available ribosomal capacity to provision the Vg2 component of an egg on its own. However, once we account for discontinuous translation and the range of potential elongation rates, *Varroa* would need to use 16% to 27% of their available ribosomal capacity for Vg production alone.

**Figure 1.**
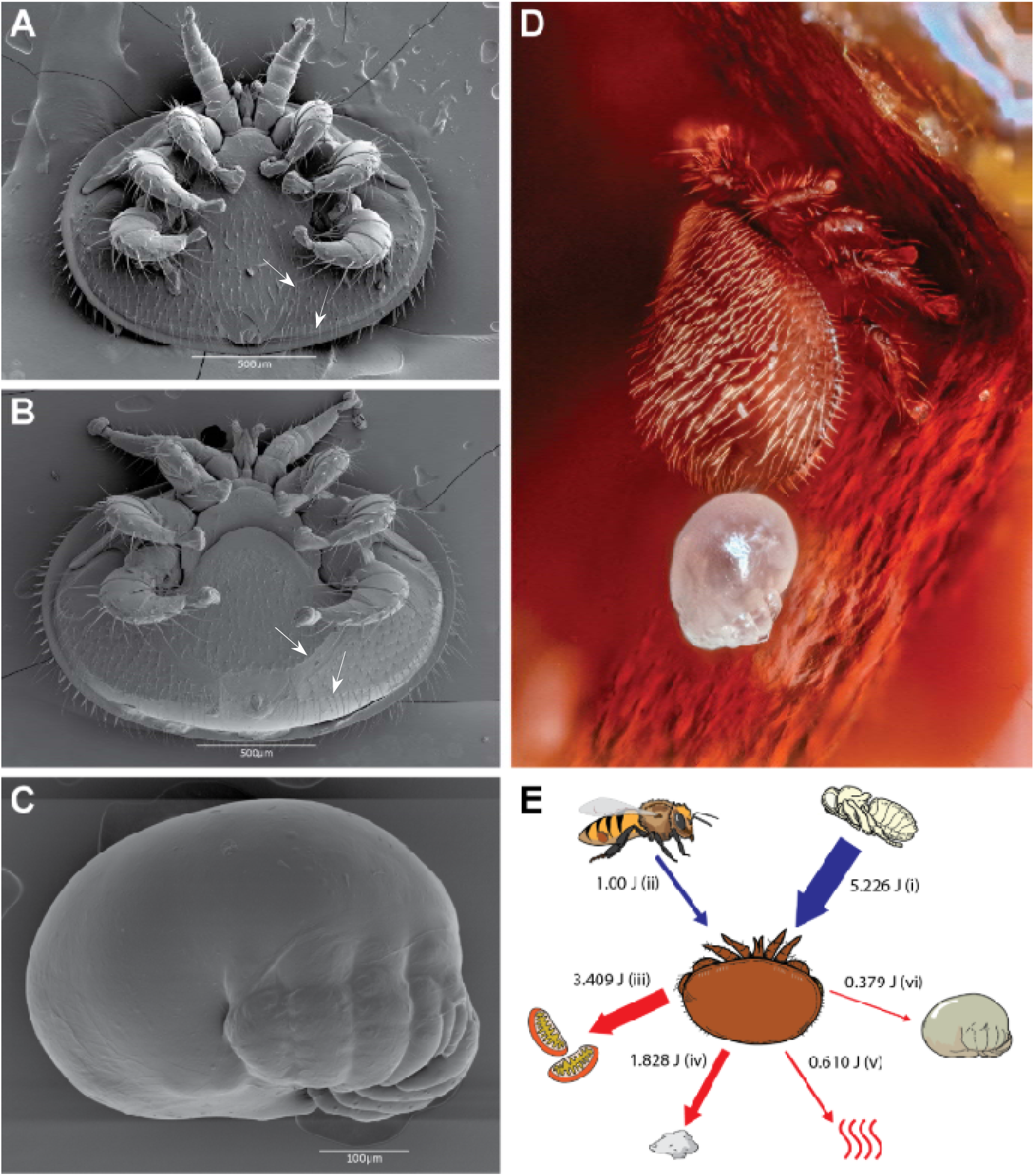
External morphology, scale, and evidence of accelerated embryonic development in *Varroa destructor* and its egg. (*A-C*) Electron micrographs detailing external ventral morphology of a non-gravid ***(A*)** and gravid **(*B*)** *Varroa destructor* (60X magnification), and **(*C*)** an egg (200X magnification). **(*D*)** Optical micrograph of a *Varroa destructor* foundress alongside a recently-laid egg inside of an *Apis mellifera* brood cell. A single egg comprises 18% of a gravid mite’s body volume (note that a gravid mite will lay one egg roughly every 30 hours). When gravid, the female’s body is substantially distended by the size of the egg, pushing all integument plates away from each other revealing a taut intersegmental membrane that is otherwise hidden (white arrows) **(*B*)**. In addition to its large size, the egg houses a fully differentiated and developed eight-legged protonymph, as opposed to a six-legged larva typical of other Mesostigmata. Features such as the legs and discrete setae are visible through the outer chorion **(*C*)**, consistent with the accelerated embryogenesis required to match the host bee’s pupation timeline. **(*E*)** Schematic diagram of the static energy budget model. Energetic inputs (blue arrows) and outputs (red arrows) within the 30-hour gestational period for an egg are shown. Energetic values for each component in the energy budget were determined with bomb calorimetry. Inputs include immature bee fat body-hemolymph aggregate (i) and stored energy from the mite’s dispersal phase (ii), during which the mite feeds on adult fat body. Energetic outputs include somatic maintenance (iii), excreta (iv), heat (v), and reproduction (vi). Despite the accelerated development within the egg, as well as its large size relative to the foundress’ body mass, the egg constitutes the lowest investment of the foundress’ total energy budget during the gestation period. Relatively low energy investment in reproduction can indicate efficient nutrient processing, but the large volume of undigested host protein we detected in the mite’s excreta suggests inefficient nutrient processing.

### 3.2 HPLC-MS/MS and immunodetection reveal *Apis* proteins in *Varroa* and *Varroa* eggs

We detected 2,287 proteins from SDS-PAGE protein gels followed by HPLC-MS/MS identification of associated bands from mite egg samples. Two *Apis* proteins were detected with normalized total spectra in the 99^th^ percentile of all detected proteins in *Varroa* eggs, those being apolipophorin and vitellogenin (**Table 1**). In total, we detected 313 *Apis* proteins in mite eggs. The majority of these proteins were detected in low abundance (normalized total spectra > 50) (***SI Appendix,* Fig. S3**). Removing these low abundance proteins resulted in seven *Apis* proteins in mite eggs. These included two members of the large lipid transfer protein (LLTP) family: apolipophorin and vitellogenin; three hexamerins (Hex): Hex110, Hex70a, and Hex70b; and two heat shock proteins (HSP): HSP90 and HSP83 (**Table 2**). Three of these proteins (Vitellogenin, HSP90, and HSP83) were detected in greater abundance in the eggs than within gravid mites. Immunodetection for *Apis* Vg confirmed the presence of this protein in both gravid *Varroa* and their eggs (***SI Appendix,* Fig. S4**). A protein fragment of approximately 40 kDa also reacted to our Vg antibodies in fat body–hemolymph aggregate from immature bees, likely a post-translationally cleaved product of the original Vg protein. This fragment was not detected via immunodetection in *Varroa* eggs.

**Table 1.**
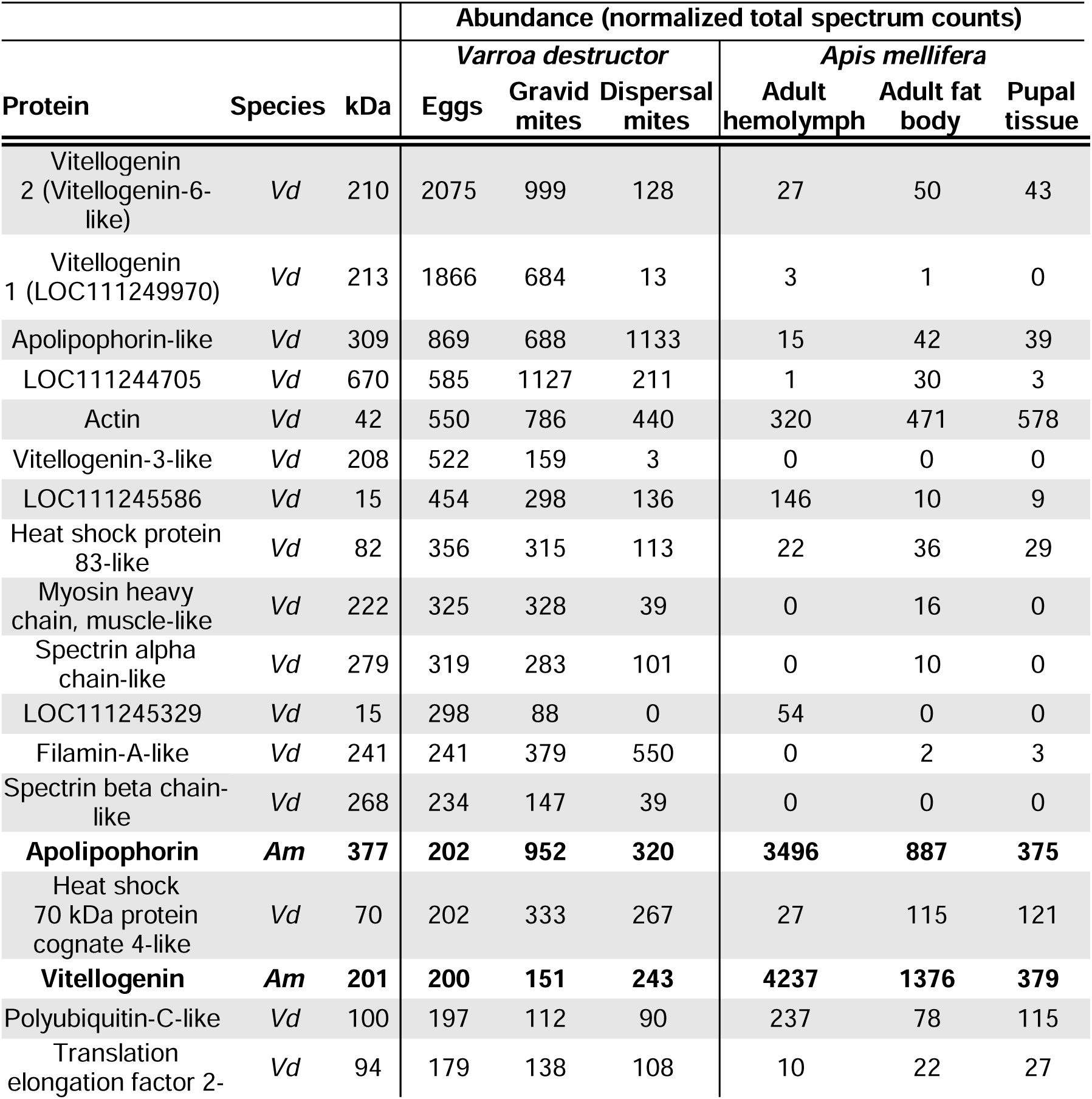
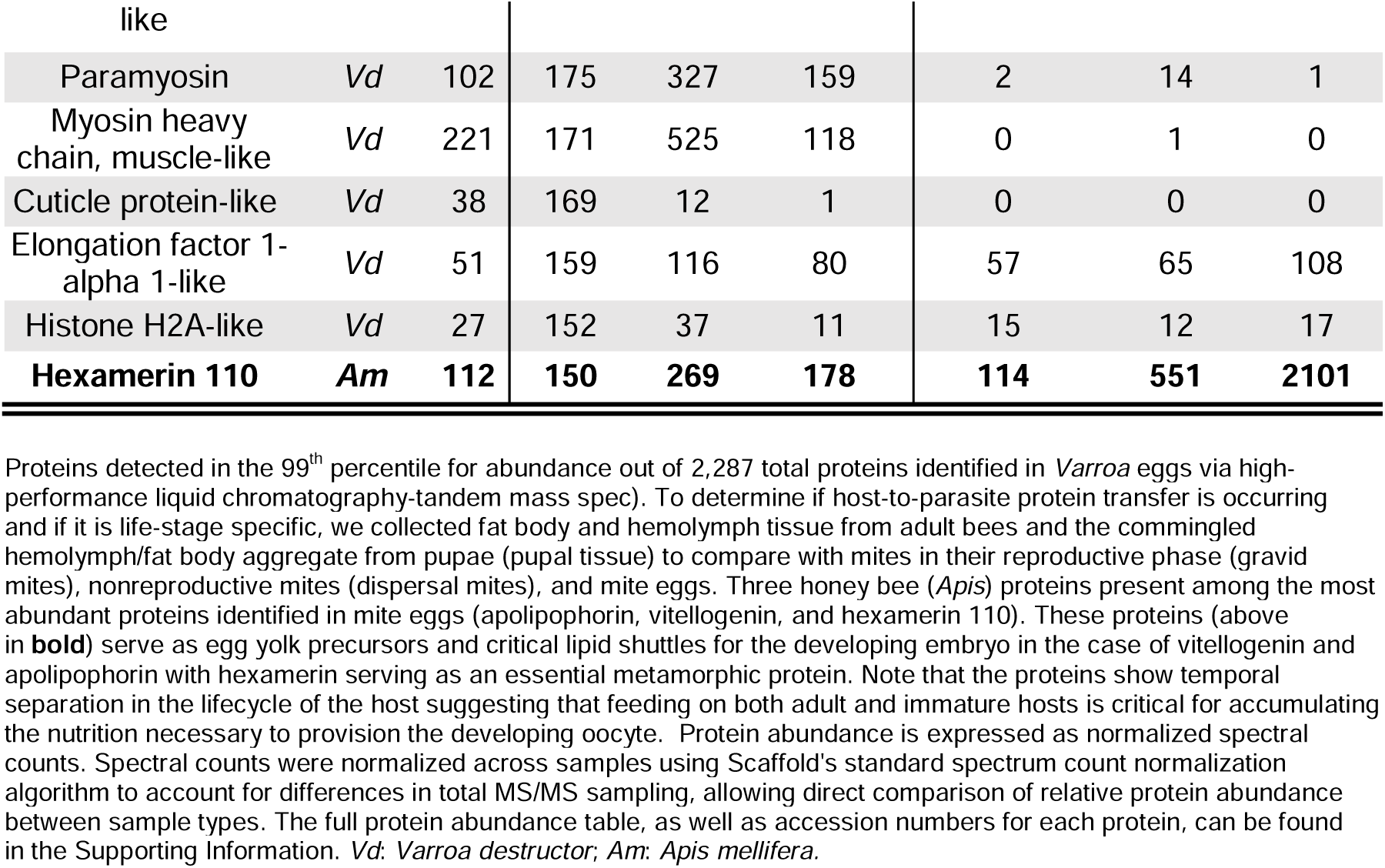
Abundant exogenous and endogenous proteins detected in Varroa eggs.

### 3.3 The lyrate organ functions as a possible site for conveyance of host proteins to ova

To better assess how the mites may be moving host proteins across the multiple membranes required to reach the egg, we used microCT to determine whether any tissues were physically connected to the ovum. We observed a dual-lobed organ, the lyrate organ, forming a connective tube with the ovum, confirming the findings of Dittman and Steiner (1997) (**Fig. 2*C-E***). This connective tube arises from the left lobe of the lyrate organ (observing the mite from the coronal plane). This tube forms a large, seemingly bundled mass at the connection point in mature ova. These scans show that the lyrate organ is a particularly dense tissue with a density similar to many of the mite’s muscles. A lyrate-ovary syncytium is visible (**Fig. 2*F and G***), which is consistent with previous work that identified an array of microfilaments between these two regions (51). This further suggests an avenue for transport from the lyrate organ into the developing ovum.

To identify the pathways facilitating the movement of *Apis-*derived proteins into the mite egg, we expressed a recombinant Vg-HaloTag construct labeled with a JF549 fluorophore and introduced it to reproductive foundress *Varroa* to track its post-ingestion localization. While several mites showed no introduced fluorescence–likely because they did not feed (***SI Appendix,* Fig. S5**)–some mites showed localized internal fluorescence. Intensity was highest in the region of the lyrate organ; the resulting fluorescence delineated the organ’s morphology, indicating that this organ accumulates *Apis* Vg (**Fig. 2*A and C***). One specimen (**Fig. 2*A* (ii), 2*B***) also showed localized fluorescence in the lower digestive tract, supporting previous observations of low proteolytic enzyme activity in *Varroa,* allowing large proteins to remain intact within the gut (26).

**Figure 2.**
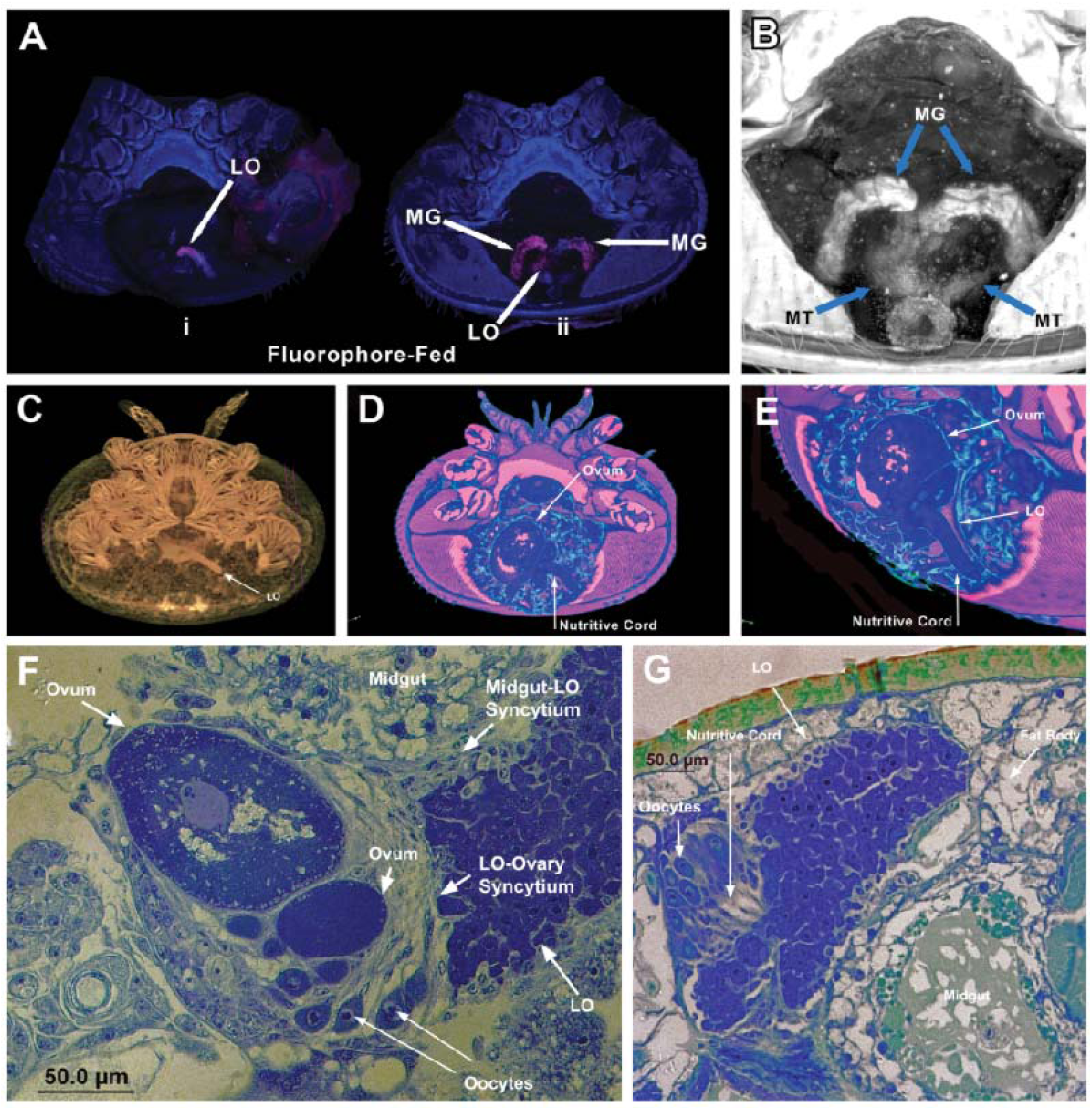
Fluorescence microscopy, X-ray computed tomography, and histology mapping molecular transport in the *Varroa* kleptocytosis pathway. (A-B) Confocal Fluorescence Microscopy: Ventral profiles of *Varroa* allowed to feed on homogenized host tissue with a Vg-fluorophore complex (Vg-HaloTag-JF549) ad libitum for 48 hours (N=8, see *SI Appendix* Fig. S5 for all mites imaged). The genital plate was removed from all specimens prior to imaging. (*A*) Though several mites did not apparently feed, mites that fed on homogenized bee tissue mixed with recombinant Vitellogenin-HaloTag-JF549 showed localized fluorescence in the dual-lobed lyrate organ (LO) (i) and the posterior lobes of the midgut gastric caeca (MG) (ii), suggesting that exogenous proteins in the parasites’ diet are conveyed to the developing oocyte via the lyrate organ allowing them to circumvent receptor-mediated uptake. The magenta channel denotes the internalized Vg-HaloTag-JF549 fluorophore signal, while the blue channel maps baseline cuticle and exoskeleton autofluorescence, determined using an algorithm to identify per-pixel correlations to fluorophore-like spectral properties (*SI Appendix* S8). (*B*) Maximum intensity projection of the TRITC channel of (*A*)(ii), with gamma adjusted to 0.47 to optimize structural visibility of internal organs. The lyrate organ sits behind the lower gastric caeca of the midgut. The proximal ends of the Malpighian tubules (MT) are visible where they empty into the post colon just superior to the anal opening. (*C-E*) X-ray Computed Tomography (CT): Regional and whole-body scans of female *Varroa* showing internal anatomy and organ structure. (*C*) Coronal whole-body computerized axial tomography (CAT) scan highlighting the location of the LO providing additional context for the tissue observed in the fluorophore-fed localizations. (*D*) Ventral view of a gravid foundress showing the nutritive cord and its continuity with the developing ovum. In the higher-magnification image (*E*), the mite is rotated slightly revealing the origin of the nutritive cord in the dense lyrate organ. (*F-G*) Tissue Histology: Light micrographs highlighting cellular junctions and nutritive infrastructure. (*F*) Coronal photomicrograph of the opisthosomal body region in a fed female mite (toluidine blue stain), showing the reproductive organs (scale bar: 50 µm). The lyrate organ (LO) forms a syncytium with the midgut and the ovary. (*G*) Sagittal photomicrograph of the opisthosomal body region (toluidine blue stain) showing nurse cells extending from the lyrate organ forming a nascent connection to early stage developing ova (scale bar: 50 µm). (*C, F, G*) Original micrographs reproduced from Sonenshine *et al* (2022) (DOI: <u>10.1093/aesa/saab043</u>) (79) and re-labeled by the authors with permission from *Annals of the Entomological Society of America*.

### 3.4 Hexamerins 110, 70b, and 70c exhibit a greater decrease in quantity than other *Apis* proteins during the deutonymph-to-adult transition

We tested the hypothesis that *Apis* hexamerins would be preferentially consumed by mites during calyptostase. Using label-free quantification (LFQ), we compared the abundances of hexamerin within deutonymphs, calyptostase mites, and teneral (newly molted) adults. Log_2_ fold-changes in LFQ intensities for each protein, relative to their abundance in the previous stage, were deemed significant if the Benjamini-Hochberg adjusted *P*-value was below 0.05. Relative to their abundances in deutonymphs, 95 *Apis* proteins in teneral adults decreased significantly, and one protein increased significantly. Relative to their abundances in calyptostase mites, no proteins significantly decreased in teneral adults, though one protein significantly increased. Relative to their abundances in deutonymphs, 44 proteins significantly decreased in calyptostases, while three proteins significantly increased. Hexamerins 70b, 70c, and 110 all decreased significantly in teneral adults, relative to their respective abundances in deutonymphs (70b: LFC = -2.81, *P* = 0.042; 70c: -3.65, *P* = 0.010; 110: -4.19, *P* = 0.023) (*SI Appendix,* Table S4, Fig. 3D). Log_2_ fold-changes from deutonymphs to calyptostases and calyptostases to teneral adults were not significant for all hexamerins, except for Hex70c, which also decreased significantly from deutonymphs to calyptostases (LFC = -1.77, *P* = 0.040). Hex110 demonstrated the largest decrease of the four hexamerins; of all 209 *Apis* proteins observed to decrease from deutonymphs to teneral adults, Hex110 was in the 94.8^th^ percentile (Fig. 3G *and H*). Compared to the other proteins, decreases in the three hexamerins most closely associated with metamorphosis (Hex110, 70b, and 70c) were greater than that of other *Apis* proteins over the entire deutonymph to teneral transition (permutation test; 100,000 permutations; observed difference = -1.400; *p* = 0.0435) (Fig. 3*A*). However, we did not observe significant differences in the transitions to and from the intermediate calyptostase (Fig. 3B *and C*) (*SI Appendix,* Table S5)

**Figure 3.**
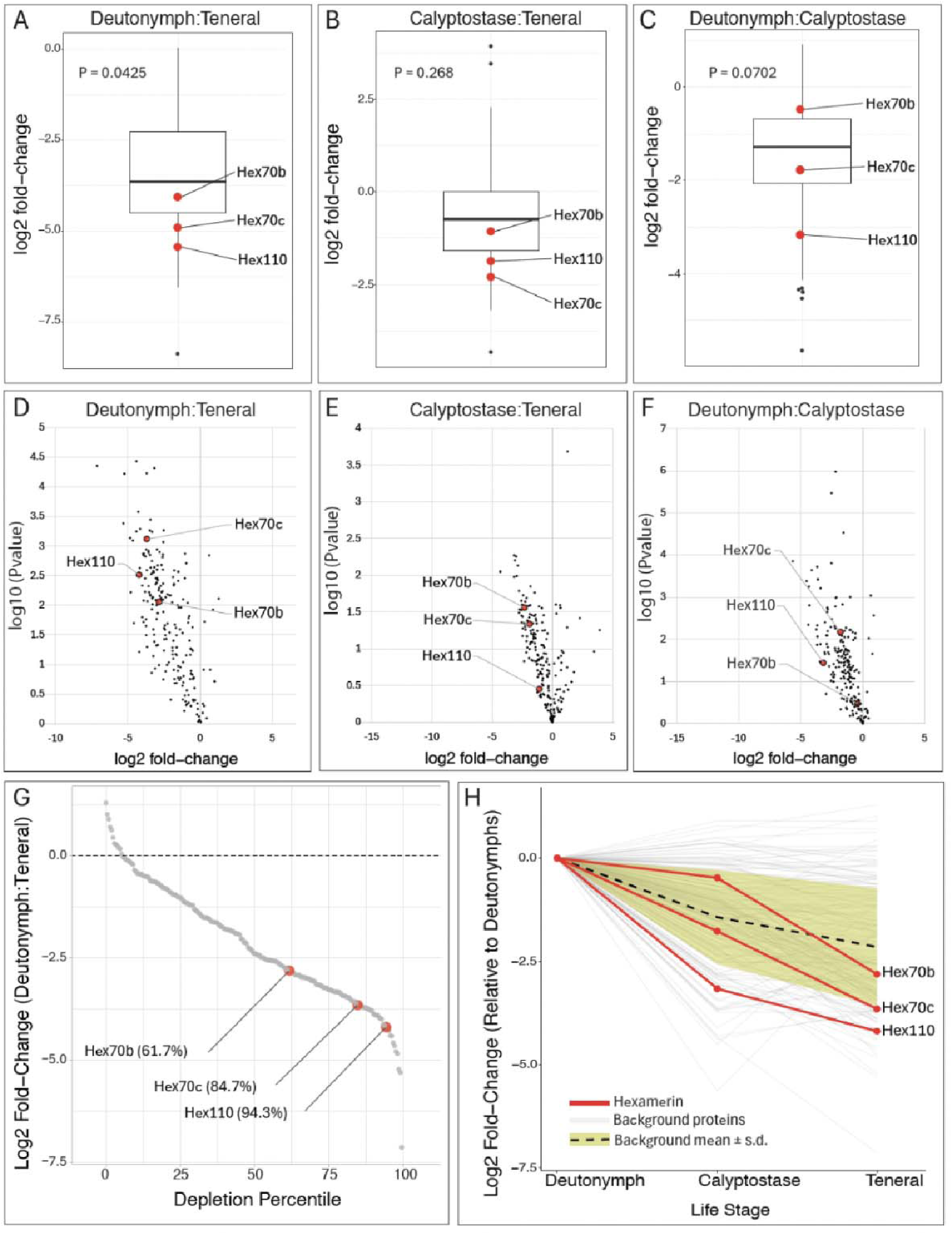
*Apis* metamorphic proteins show greater quantitative during periods of rapid reorganization relative to other ingested *Apis* proteins. Log2 fold-changes in LFQ intensities of 209 *Apis* proteins, detected through mass spectrometry, were determined in a pairwise manner between three discrete stages of *Varroa* mite metamorphosis: deutonymphs (immediately prior to metamorphosis, n = 4), calyptostase (during metamorphosis, n = 4), and teneral adults (immediately after, n = 4). All *Apis* proteins were expected to decrease during this period of rapid reorganization due to digestion by the parasite, however the three hexamerins most strongly associated with metamorphosis in insects (Hex 110, Hex 70b, and Hex 70c) demonstrated a significantly greater mean decrease over this transitional period relative to all other “background” proteins suggesting that stolen host proteins may retain their functionality in the parasite. (*A-C*) Boxplots of LFCs in all *Apis* proteins for each pairwise comparison between *Varroa* life stages. Hexamerins associated with metamorphosis are highlighted in red. P-values represent the result of the permutation test (one-tailed, nperm = 100,000), comparing mean LFC of the background proteins to that of the highlighted hexamerins (Deutonymph: Teneral, mean difference = -1.400, *P* = 0.0425; Calyptostase: Teneral, mean difference = -0.374, *P* = 0.268; Deutonymph: Calyptostase, mean difference = **-**1.000, *P* = 0.0702). (*D-F*) Volcano plots of *Apis* proteins of each pairwise comparison between *Varroa* life stages, with hexamerins highlighted in red. (*G*) Waterfall plot comparing the percentiles of LFC values for each detected *Apis* protein between deutonymphs and teneral adults. Hexamerins (with percentiles) are highlighted in red. Note that hexamerin 110 demonstrates the greatest decrease amongst other hexamerins and is in the 94^th^ percentile of all LFC over the entire studied developmental period. (*H*) LFC for every *Apis* protein, relative to deutonymphs, showing trends over time to the teneral adult stage. Hexamerins are highlighted in red, background proteins are in grey. The mean ± s.d for the background *Apis* proteins are overlayed (black dashed-line and yellow bands, respectively).

**Table 2.** Most abundant *Apis*-derived proteins detected in *Varroa* eggs.

| Protein | Species | kDa | Abundance (normalized total spectrum counts) |  |  |  |  |  |
| --- | --- | --- | --- | --- | --- | --- | --- | --- |
|  |  |  | *Varroa destructor* |  |  | *Apis mellifera* |  |  |
|  |  |  | Eggs | Gravid mites | Dispersal mites | Adult hemolymph | Adult fat body | Pupal tissue |
| Apolipoporphin | *Am* | 377 | 202 | 952 | 320 | 3496 | 887 | 375 |
| Vitellogenin | *Am* | 201 | 200 | 151 | 243 | 4237 | 1376 | 379 |
| Hexamerin 110 | *Am* | 112 | 150 | 269 | 178 | 114 | 551 | 2101 |
| Heat shock protein 90 | *Am* | 83 | 128 | 91 | 33 | 31 | 110 | 100 |
| Hexamerin 70a | *Am* | 81 | 88 | 150 | 92 | 21 | 78 | 329 |
| Heat shock protein 83 | *Am* | 83 | 77 | 62 | 24 | 11 | 41 | 52 |
| Hexamerin 70b | *Am* | 80 | 54 | 308 | 59 | 13 | 60 | 238 |

*Apis* proteins found in mite eggs in high abundance, as determined through high-performance liquid chromatography-tandem mass spectrometry. Total spectrum counts were normalized across samples using Scaffold's standard spectrum count normalization algorithm. Proteins were classified as high abundance if the normalized total spectrum counts were greater than 50. Of the total 313 detected *Apis* proteins in *Varroa* eggs, seven were observed in high abundance. Notably, these high abundance proteins represent crucial nutritional and developmental resources for embryonic growth. Additionally, vitellogenin, heat shock protein 90 and heat shock protein 83 are present in higher abundance in the eggs than in the gravid mites suggesting that the *Varroa* foundress is selectively provisioning the egg with host proteins to fuel rapid embryonic development and oogenesis. The presence of proteins from a variety of different protein families is consistent with the conclusion that *Varroa* have evolved a means of bypassing receptor-mediated uptake to convey nutrients to developing ova. The full table of *Apis* proteins in mite eggs, as well as accession numbers for each protein, can be found in the Supporting Information. *Am*: *Apis mellifera*.

## 4. Discussion

Kleptoparasitism is defined as the intra- or interspecific usurpation of a resource that another individual has invested energy into procuring or producing (52, 53). Our work demonstrates that *Varroa* steal and transfer relatively large, complex proteins into their ova from immature and adult honey bees. We call this process “kleptocytosis,” denoting the movement of stolen and intact molecules into a cell. These proteins are not digested and repurposed as in normal parasitic feeding processes but apparently serve the same or similar purpose as in the host organism making this a distinct adaptation from typical nutrient acquisition. The selective uptake of these proteins, transport of these proteins, unique anatomical adaptations of the mite, and apparent transport from the gut into the lyrate organ suggest that this system is of critical importance to the parasite.

Our static energy budget revealed that *Varroa* invest less energy in reproduction than energy lost to respiration or any other energetic context contributing only 9.99% of their available energy to the process. This investment in reproduction is dwarfed even by the energy in their waste with 4.8 times more energy in their excrement than the energy incorporated into the egg. The energy demand associated with rapid reproduction is comparatively high but our calculations show that the foundress’s reproductive output is limited not by caloric intake but by the mite’s capacity to convert energy into specific macromolecular constituents required for the rapid development of the oocyte. To be capable of meeting the energetic demand of producing large, well-developed offspring, *Varroa* would need to be highly efficient at processing nutrients but previous study has shown that *Varroa* are markedly *inefficient* at nutrient processing (34).

Despite the low reproductive investment, *Varroa* are capable of meeting the energetic demand of producing large, well-developed offspring in 30 hrs; with each egg approximately 18% of the foundress’s body volume and 14% of its body mass *(**SI Appendix,*** **Table S3**). Additionally, there is enough energy invested in each egg to support the embryo’s post-ovipositional accelerated development, such that it hatches within 24-30 hours as an eight-legged protonymph. (54). For comparison, Triatomine kissing bugs allocate 68% of their adult energy income towards reproduction and produce an average reproductive investment comprising 12% of the adult body mass (55) (***SI Appendix,* Table S6**). For a nearly seven-fold lower relative energy investment, *Varroa*’s reproductive investment is similarly proportional by mass to that of a kissing bug.

While *Varroa* carry multiple ova in development at once (**Fig 2*F***), only the most mature ovum is provisioned at a time, inhibiting the development of later ova until the most mature egg is laid (56). This limits the time that the foundress has to produce the full complement of proteins needed to provision the developing ovum to 30 hours, representing a translational challenge. By our estimate, under realistic conditions, a *Varroa* foundress would need to use 16 to 27% of its total ribosomal capacity to produce enough Vg to provision a single ovum. However, of the foundress’ total ribosomal capacity, it is likely that at least 50% of ribosomes are encumbered to produce housekeeping proteins, such as ribosome components (45). When we include ribosomes that are available to produce non-essential proteins, we estimate Vg synthesis to require between 32% and 54% of these ribosomes. While Vg is by far the most abundant protein in acarine ova (35,36), we identified more than 2,000 other proteins which would require additional transcripts and dedicated time for translation. This represents a substantial challenge which the foundress cannot meet by slowing reproduction as would be possible in species not subject to their host’s developmental timeline.

The capacity to surmount such a reproductive challenge without developing corresponding nutrient processing efficiency requires biochemical explanation. Through selective uptake of host egg yolk precursors and metamorphic proteins, kleptocytosis would relieve the biosynthesis bottleneck allowing mites to successfully complete their lifecycle before their window provided by their metamorphosing host closes.

While we detected 313 *Apis* proteins inside *Varroa* eggs, the majority of these were in low abundance, especially within the context of the global *Varroa* egg proteome. However, three *Apis* proteins were observed in high abundance, such that they were in the 99^th^ percentile of all proteins within mite eggs. These proteins were apolipophorin, vitellogenin, and hexamerin 110. The presence and high quantity of these energy-dense proteins support our hypothesis that *Varroa* are selectively extracting nutritionally complex proteins for the developing oocyte, aiding accelerated development within the egg. Vitellogenin in particular is an energy-rich egg-yolk precursor and serves as the primary nutrient donor for the developing oocyte (57). While mites can synthesize their own vitellogenins (23), kleptocytosis may have developed as an adaptation to ensure the mite’s life cycle is completed during the brief time frame in which their host is pupating. Additional vitellogenin which does not need to be synthesized would ostensibly allow the parasite to produce eggs more quickly. Because *Varroa* offspring die if they are not fully mature by host eclosion (2), the time saved by forgoing the *de novo* synthesis of essential reproductive and developmental proteins provides a critical fitness advantage. In addition to Vgs, apolipophorins serve as the primary egg-yolk precursor for other arthropod species, and, along with other LLTPs, are heavily involved in lipid transport during embryonic development (58, 59).

Hexamerins are another energy-rich protein family that is primarily involved in insect metamorphosis. Hexamerins are a family of storage proteins which expand the amino acid pool needed to complete metamorphosis during the pupal stage, during which the insect cannot feed and incorporate additional amino acids from the diet (19, 60). Notably, arachnids do not produce hexamerins (though hemocyanins may function as an analog in some species) (61). While *Varroa* lack a pupal stage, they instead have defined calyptostatic stages in their lifecycle, which are analogous to insect pupal stages (16). This is unique amongst other mites in the order Mesostigmata, which do not go through calyptostase. One of these calyptostatic stages occurs within *Varroa* eggs, during which the larva develops a fourth pair of legs in less than a day, ostensibly requiring a substantial biochemical investment of storage proteins not present in the mite’s proteome. Hexamerins, transferred from the host via kleptocytosis, may provide the energy and amino acid pool needed to support the larval metamorphosis within the egg.

As hexamerins are metabolized to fuel metamorphosis, we expected that the rate of host hexamerin depletion would significantly exceed that of non-storage host background proteins during calyptostase. While the transition between larval *Varroa* and protonymphs would have been the preferred system for studying hexamerin usage during calyptostase, there is no clear “before” stage for comparison. As such, we focused our attention on the deutonymph to adult transition, assuming the processes involved during the calyptostatic stage between these two stages would be similar to the process that occurs in the larva-protonymph transition within the egg. We observed significant decreases in all four *Apis* hexamerins (70a, 70b, 70c and 110) over the entire deutonymph to teneral transition (***SI Appendix,* Table S4**), as well as a dramatic log_2_ fold-change for Hex110 (placing it just under the 95^th^ percentile of most decreased host proteins between deutonymphs and adults). In addition, when focusing on hexamerin 110, 70b, and 70c, we observed a mean log fold-change more negative than that of the other *Apis* proteins, a pattern consistent with the function and temporal trends of hexamerins in their original host. We excluded hexamerin 70a from this analysis because of its unique depletion pattern relative to the other hexamerins. In developing honey bees, Hex110, 70b, and 70c are depleted rapidly during and soon after metamorphosis (62). In contrast, Hex70a abundances remain elevated relative to other hexamerins several days after emerging as an adult (62). This is likely due to pleiotropic functioning of Hex70a. In addition to acting as a reservoir for metamorphosis, hex70a is crucial for gonad development (63). We observed similar trends for *Apis* Hex70a in *Varroa*; between deutonymphs and teneral adults, Hex70a decreased the least of the four hexamerins. Together, these data support our hypothesis that host hexamerins are utilized by *Varroa* to aid their rapid development.

The presence of both *Apis* Vg and hexamerins within the egg may also explain the need for *Varroa* to feed on both adult and immature bees to consistently initiate reproduction (14), as there is a temporal divide in the mite’s access to these proteins. While feeding on brood allows the mites to feed on a defenseless host inside a protected cell, feeding on adult bees exposes mites to grooming and the threat of falling off a host and out of the colony. Thus, there would have to be a strong evolutionary benefit to counteract this fitness drawback. Xie *et al.* (2016) demonstrated that foundress mites have lower reproductive success when prevented from feeding on nurse-aged bees, speculating that there may be a nutritional underpinning for this (14). Our finding of *Apis* vitellogenin in the mite eggs provides this nutritional link, as Vg is produced primarily in adult bees, with nurse-aged bees carrying the highest levels (64–66). Notably, mites have lower, though non-zero, reproductive success when not feeding on adult bees (14). Given that *Varroa* lose access to nurse bees while confined to a brood cell, we expect that reproductive mites would have to draw on a reservoir of Vg stored from their time feeding on adult bees. These temporal patterns of vitellogenin are in contrast to hexamerins, which are most abundant in immature bees (66).

In addition to the LLTPs and hexamerins, we also observed an assemblage of other host proteins including the heat shock proteins HSP90 and HSP83, myosin heavy chain, and the LLTP larval-specific very high-density lipoprotein, though at lower abundances than apolipophorin, Vg, and the hexamerins (**Table 2**). These proteins are interesting for their reproductive utility. Both HSP83 and HSP90 are chaperone proteins which are necessary for embryogenesis and oogenesis in diverse clades of insects (67–70). The roles of myosin heavy chain proteins in reproduction are not well understood, but they have been shown to be essential for insect embryonic development (71). We also detected *Varroa* proteins within *Apis* samples, likely indicating a bidirectional molecular exchange at the feeding site. High-abundance proteins such as *Varroa* actin and even *Varroa* HSPs likely represent salivary products injected to aid the mite’s feeding and extraoral digestion (72–75), or backflow from *Varroa* into *Apis* samples during the feeding process. We distinguish this incidental crossover from kleptocytosis based on the asymmetry of nutritional protein abundance and selective sequestration in reproductive tissues by the mite.

In addition to the three most abundant *Apis* proteins in *Varroa* eggs, we also observed 313 other proteins, though most were at low abundance. The diverse assemblage of host proteins found in *Varroa* eggs suggests an alternative pathway for *Apis* proteins to enter the egg from the gut, bypassing the normal means for provisioning the developing ovum. Developing ova are biochemically closed systems; components are introduced to the oocyte via highly specific receptor-mediated transport. For example, receptors for vitellogenin (Vg), the primary egg-yolk precursor in most insects, are specific enough to exclude Vgs from closely related species (22, 31). Given the evolutionary distance between *Apis* and *Varroa*, it is unlikely that *Varroa* proteins and those of its host would be able to use the same receptor, raising the fundamental question of how *Varroa* mites could provision their ova with exogenous proteins (76, 77).

Our data suggest that the lyrate organ is the likely pathway for *Apis* proteins to be introduced into the developing egg, as well as serving as the storage site for *Apis* proteins that are acquired while feeding on adult bees. In other mite species, the lyrate organ is a component of the reproductive system, providing the ovaries with proteins, lipoproteins, and other nutrients (78). This organ forms a cytoplasmically continuous syncytium with the midgut epithelium, allowing for host nutrients to flow seamlessly from the digestive system into the lyrate organ (**Fig. 2*F and G***) (51, 79). Furthermore, the lyrate organ contains numerous elongated nurse cells, which extend to the developing oocyte, forming an intercellular connection with the ovum (**Fig. 2*G***) (80). Together, this suggests a continuous pathway for undigested *Apis* proteins from the midgut to be transferred into the egg, using the lyrate organ as an intermediary. MicroCT imaging revealed that the lyrate organ remains connected to the oocyte throughout the latest stages of oocyte development, allowing a continuous stream of nutrients into the egg (**Fig. 2D and *E***).

After feeding *Varroa* our Vg-HaloTag-JF549 construct, we observed a fluorescent signal localized to an area that appeared to match the lyrate organ in morphology and locality. The mites used in this study were not gravid, suggesting that host vitellogenins, and likely other proteins, accumulate in the lyrate organ prior to transport into the developing oocyte (**Fig. 2**). While we observed some fluorophore-like signal from the control mites, this signal was unstructured and unbound by defined organs compared to the signal identified in mites which were fed the fluorophore-vitellogenin construct, indicating that the control signal was likely from the mite’s autofluorescence. We observed fluorophore signals from only two of the eight mites exposed to the vitellogenin-fluorophore construct; this may be due to unequal ingestion of the fluorophore between mites (***SI Appendix,* Fig. S5**). Additional fluorescent microscopy feeding trials are necessary to confirm that this pathway is consistent across exogenous proteins. Given that Vg, HSP90, and HSP83 were detected in higher concentration in the eggs than the foundresses, they are likely being trafficked via an active transport channel that may differ from other host-derived proteins.

Digesting and synthesizing complex reproductive/metamorphic proteins is an expensive and time-consuming process. Stealing them from the host represents an adaptation that enables the creation of developmentally advanced offspring with a relatively low energy investment. This is especially critical in the sensitive context of the *Apis–Varroa* host–parasite complex in which the parasite must develop on a precise timeline or lose its chance at reproduction. As such, study of this pathway provides valuable insights into the biomolecular plasticity of metamorphosis, invertebrate embryonic development, and a broader understanding of alternative reproductive strategies in parasites. Additionally, kleptocytosis is likely one of the key adaptations that has allowed this parasite to exploit a well-defended host with few other metazoan parasites. Understanding this pathway reveals a potential vulnerability in a parasite that has proven to be exceptionally robust. Our work suggests that *Varroa* need these host proteins (some of which they are unable to synthesize themselves) in order to complete their life cycle. In the context of *Varroa*’s strict developmental and reproductive timeline, a short delay in these processes reduces the parasite’s reproductive fitness to zero. As such, this work suggests that future attempts to control *Varroa* do not need to fully sterilize the parasite in order to be effective. Future work will need to target understanding the individual steps in this unique pathway and identify any potential weak links that may help us disrupt the strict developmental synchrony required for reproductive success.

## Data Availability Statement

Raw data and code supporting the findings of this manuscript are available on Zenodo (10.5281/zenodo.21228128), to be made public upon publication.

## Acknowledgments

We would like to thank the USDA Agricultural Research Service’s Bee Research Laboratory for providing invaluable training and assistance throughout this project. We thank Dr. Tom Cech for providing mentorship and editorial comments on the manuscript. We are grateful to Dr. Naresh Chandra Tewarson for his painstaking research in *Varroa* biology, without which we would understand substantially less about this parasite. We thank Dr. Brenda Klaunberg and Danielle Donahue of the National Institutes of Health, Mouse Imaging Facility for allowing us to conduct our initial scans of *Varroa* in their facility. Fluorescence microscopy imaging was performed at the BioFrontiers Institute’s Advanced Light Microscopy Core (RRID: SCR_018302). The Nikon AXR Laser Scanning Confocal used in this manuscript is supported by NIH Grant 1S10OD034320. Label-free quantitative Mass spectrometry was performed at Proteomics and Mass Spectrometry Core Facility (RRID: SCR_ 018992) in the Department of Biochemistry at the University of Colorado Boulder for data acquisition and analysis using the Thermo Orbitrap Q-Exactive HF-X, funded by NIH S10-OD025267. Mention of trade names or commercial products in this publication is solely for the purpose of providing specific information and does not imply recommendation or endorsement by the USDA. The USDA is an equal opportunity provider and employer. This study was supported jointly by a grant from Project Apis m., the Central Maryland Beekeepers Association, the Santa Clara Valley Beekeepers Guild, the Eppley Foundation for Research, and the Ramsey Research Foundation.

