## Supplementary Figures for "Kleptocytosis: Comparative Proteomics and Functional Imaging Show *Varroa destructor* Co-opts Intact Host Proteins for Rapid Development and Reproduction"

**This PDF file includes:**

Supporting text

Figures S1 to S6

Tables S1 to S8

Legends for Datasets S1 to S2

SI References

**Other supporting materials for this manuscript include the following:**

Datasets S1 to S2

Supporting Information Text

**S1. Energy lost as heat**: To measure energy lost as heat, 177 gravid mites were placed into a sealed inert glass container of known volume (35.32 ml). A high precision temperature sensor (±0.1°C) (TMP117, Texas Instruments) was queried once per second after being sealed with reflective ducting tape. Once sealed, a gravid mite sample container was placed into an incubator pre-heated to 28.6°C alongside a container containing non-gravid mites, and an empty control container, each with redundant temperature sensors and logging devices to account for heat contributed by the test-harness. Data were analyzed using a [nth] polynomial fit model matching the temperature decay of each respective chamber. The temperature of the gravid mite container remained higher than that of the controls as expected. The point of first-first order convergence indicating mite-death was noted, and differences between gravid and control slopes were subtracted to yield gravid mite energy contribution before death. The resulting value, assumed to be the basal metabolic energy contribution by Varroa, was used to calculate the energy necessary to heat the container volume in a 30-hour timeframe, yielding 0.61 J per mite.

**S2. Bomb calorimetry.** Samples, kept on dry ice, were weighed on a VWR precise weighing balance (VWR-220AC) with all the transfer tools (e.g., brush and forceps), and then added directly to the cup holder. For the *Varroa* egg samples, the samples were covered in a benzoic acid spike (Parr Instrument Company, Moline, IL, Cat# 3414) to ensure complete and efficient burning. A Parr oxygen bomb semimicro calorimeter (Parr Instrument Company, Moline, IL, model number 6725) and a calorimetric thermometer (Parr Instrument Company, Moline, IL, model number 6772) were used to obtain the data. Egg samples were run with spike correction enabled, and data were calculated based on sample weight and spike weight, adjusting for the remaining fuse wire according to the manufacturer’s protocol.

**S3. Antibodies to *A. mellifera* *and* *V. destructor* vitellogenin.** Amino acid sequences for *A. mellifera* (NP_00101157) and *V.* *destructor* (AFN88463 and AFN88464) vitellogenin were examined. Comparison of pairwise alignments (MUSCLE) for the A. mellifera vitellogenin amino acid sequence (NCBI: NP_001011578, 1770 aa) versus the primary V. destructor Vg amino acid sequence (NCBI: AFN88463, 1850 aa) showed only 17.2% identical sites, and only 35.0% pairwise positive similarity (BLSM62). Amino acid regions likely to contain immunogenic sites were identified and submitted to the Thermofisher Life Sciences custom antibodies production service (Thermofisher Life Sciences, 14665 Rothgeb Dr, Rockville, MD 20850) for producing antibodies raised in rabbits. For the series of antibodies selected to sensitize the rabbit host, the sequences and target peptides are shown in **Table S7**. BLASTp matching shows 79.1% similarity with *Apis cerana*, 76.4% with Vg from *A. mellifera* (AEQ58342), but does not match with *V. destructor*.

**S4. MS and MS/MS.** MS and MS/MS was performed at the Keck Biomolecular Research Facility at the University of Virginia (Charlottesville, VA). The excised bands were digested with trypsin, and the extracted peptides were dissolved in 50% acetonitrile/5% formic acid, concentrated, and analyzed. The LC-MS system consisted of a Thermo Electron Q Exactive HF-X mass spectrometer system with an Easy Spray ion source connected to a Thermo 75 µm x 15cm C18 Easy Spray column (through pre-column). Of the extract, 5 µL of the extract was injected and the peptides eluted from the column by an acetonitrile/0.1M acetic acid gradient at a flow rate of 0.3 µL/min over 1 hour.

**S5. MicroCT methods for mite and egg volume.** Non-gravid adult mites and mite eggs were collected from beehives using small paint brushes and stored at -70ºC before imaging. Eggs were not considered viable if they had legs extending outward evincing successful eclosion from the egg stage. MicroCT images were acquired using a ZEISS Xradia 520 Versa X-Ray CT machine. Mite samples were imaged in a polymer tube and stabilized using foam. Egg samples were imaged in a polymer tube by placing them on the side of the tube via paintbrush. Sample isolation and volume calculation was done in Dragonfly 3D Visualization and Analysis software.

**S6. HaloTagged Vitellogenin Fusion Protein Expression and Purification.** Vitellogenin gene fragments were sequentially linked into a plasmid construct via Gibson assembly. This construct included a hexahistidine and maltose binding protein (MBP) fusion with a PreScission protease cleavage site on the N-terminal end of the vitellogenin coding sequence, as well as a HaloTag protein fusion on the C-terminal end.

This plasmid was used to make infectious baculovirus stock in Sf9 (*Spodoptera frugiperda*, IPLB-Sf-21-AE) cells using the Bac-to-bac system (Invitrogen). To express recombinant protein, High Five (*Trichoplusia ni*, BTI-Tn-5B1- 4) cells were transfected with baculovirus at 28°C for 66 hours, frozen in liquid nitrogen, and stored at -80°C until use. All protein purification steps were performed in a 4°C cold room. Cells were lysed in lysis buffer (30 mM Tris-HCl (pH 7.5) at 4°C, 250 mM NaCl, 2 mM MgCl_2_, 1 mM TCEP, 10 mM imidazole, 0.5% NP-40, 10% glycerol, 25 mM L-arginine, 25 mM L-glutamate, and protease inhibitor cocktail) for 1 hour and sonicated with mild strength. Debris was then removed by centrifugation at 23,000 x g for 40 minutes. The supernatant was incubated with Ni-NTA agarose resin (Qiagen) for 30 minutes, then put onto a gravity flow column and washed with 10 column volumes (CV) of lysis buffer, 10 CV of high-salt wash buffer (30 mM Tris-HCl pH 7.5 at 4°C, 1 M NaCl, 2 mM MgCl_2_, 1 mM TCEP , 0.01% NP-40, and 10% glycerol), and 20 CV of low-salt wash buffer (20 mM Tris-HCl (pH 7.5) at 4°C, 150 mM NaCl, 2 mM MgCl_2_, 1 mM TCEP , 30 mM imidazole, and 10% glycerol). Proteins were then eluted in elution buffer (20 mM Tris-HCl (pH 7.5) at 4°C, 150 mM NaCl, 2 mM MgCl_2_, 1 mM TCEP, 300 mM imidazole, 10% glycerol, 25 mM L-arginine, 25 mM L-glutamate, and protease inhibitor cocktail) and dialyzed in 1 L buffer (20 mM Tris-HCl (pH 7.5) at 4°C, 300 mM NaCl, 1 mM TCEP , and 10 mM EDTA) for 2-3 hours to remove imidazole, then in 1 L amylose column buffer overnight (20 mM Tris-HCl (pH 7.5) at 4°C, 250 mM NaCl, 2 mM TCEP, and 2 mM EDTA).

The dialyzed solution was then incubated with Amylose resin (Qiagen) for 30 minutes, and resin was washed with 10 CV of high-salt wash buffer (10 mM Tris-HCl pH 7.5 at 4°C, 500 mM NaCl, 2 mM MgCl_2_, and 50 mM EDTA), and 20 CV of low-salt wash buffer. Proteins were then eluted in elution buffer (10 mM Tris-HCl (pH 7.5) at 4°C, 300 mM NaCl, 2 mM MgCl_2_, 1 mM TCEP, 50 mM EDTA, and 10 mM maltose). 1 mg of PreScission protease per 50 mg of protein was then added to the eluent and left at 4°C overnight.

An AKTA-FPLC system was then used for subsequent purification with a Superose 6 increase 10/300 column (GE Healthcare). The Superose 6 increase 10/300 column was equilibrated and performed with buffer (20 mM Tris-HCl (pH 7.5) at 4°C, 250 mM NaCl, 1 mM TCEP, 2 mM EDTA, and 10% glycerol). The protein complex was flash-frozen in liquid nitrogen as single-use aliquots and stored at −80°C.

**S7. Fluorophore attachment.** To attach the fluorophore, single-use 0.5 mL aliquots of the HaloTagged vitellogenin were brought up to 1 mL with buffer (2x PBS, 10% glycerol, 1 mM EDTA). Next, 5 µL of fluorophore (JF549, Howard Hughes Medical Institute Janelia campus, Ashburn, VA) was added and incubated in a photo-proof tube at 4°C for 1 hour. The solution was then added to a 100 kDa MWCO spin column, and centrifuged at 14,000g for 4 minutes to remove free fluorophore. After addition of 0.5 ml of buffer (2x PBS, 2% glycerol, and 1 mM EDTA) to the spin column it was centrifuged at 14,000g for 4 minutes. This step was repeated. The resulting solution was transferred to a new 100 kDa MWCO spin column. Another 0.5 ml of buffer was added and the sample was spun again at 14,000g for 4 minutes. Then, 0.5 mL of a second buffer (2x PBS and 1 mM EDTA) was added and spun again at 14,000 g for 4 minutes. This step was repeated twice. Finally, 0.5 mL the second buffer (2x PBS and 1 mM EDTA) was added, and spun again at 14,000g until the solution was concentrated to about 50 µL. The sample was then stored on ice.

**S8. Image analysis algorithm.** Microscopy images with fluorophore-specific signal overlay were generated in R v4.3.1 via a custom script, which has been uploaded to Github (https://github.com/whemphil/MiteImageProcessingAlgorithm). Briefly, TRITC-, EGFP-, and AF647-channel image sets were loaded into R and decomposed into signal matrices, reprocessed into a single, filtered matrix using a custom algorithm, then edited for contrast, overlayed with the original TRITC-channel image matrix, and exported as dichromatic images.

Specifically, the algorithm can be formulated as Eq. S1,

(Eq. S1)

$\overline{F}= \overline{T} \left( \overline{{R^{\circ}}} \right)^{1.8} \left( \overline{C} \right)^{5}$

where *F* is the filtered image matrix produced by the algorithm, *T* is the original TRITC-channel image matrix, *R°* is a 0-1 scaling of the matrix *R* given by Eq. S2.1, which reduces signal in pixels with spectral properties resembling mite exoskeleton autofluorescence, and *C* is a matrix given by Eq. S2.2, which expresses per-pixel correlation to fluorophore-like spectral properties. All arithmetic operations between matrices are pairwise.

(Eq. S2.1)

$\overline{R}= -\overline{N_{x}}\cos\left( -\tan^{-1} \left( -1.31 \right) \right)+ \overline{N_{y}}\sin\left( -\tan^{-1} \left( -1.31 \right) \right)$

(Eq. S2.2)

$$\overline{C}= \left[ \left( \frac{\left( \overline{N_{x}} +1.33 \right) \cos\left( -\tan^{-1} \left( -0.127 \right) \right) + \left( \overline{N_{y}} - 0.137 \right) \sin\left( -\tan^{-1} \left( -0.127 \right) \right)}{4} \right)^{2}+ \left( \left( \overline{N_{x}} +1.33 \right) \sin\left( -\tan^{-1} \left( -0.127 \right) \right) - \left( \overline{N_{y}} - 0.137 \right) \cos\left( -\tan^{-1} \left( -0.127 \right) \right) \right)^{2}+1 \right]^{-1}$$

In Eq. S2, *N_x_* and *N_y_* are normalized matrices given by Eq. S3.1 and S3.2, respectively,

(Eq. S3.1)

$$\overline{N_{x}} = {log}_{2}\left( \frac{\overline{A} t_{med}}{\overline{T} a_{med}} \right)$$

(Eq. S3.2)

$$\overline{N_{y}} = {log}_{2}\left( \frac{\overline{E} t_{med}}{\overline{T} e_{med}} \right)$$

where *A*, *E*, and *T* are respectively the original AF647-, EGFP-, and TRITC-channel image matrices, *a_med_*, *e_med_*, and *t_med_* are the median values of those matrices, and all arithmetic operations between matrices are pairwise.

**S9. Label Free Quantitative Mass Spectrometry**. Varroa mites were homogenized in 5% (w/v) SDS, 100 mM Tris-HCl, pH 8.5, 10 mM tris(2-carboxyethylphosphine) (TCEP) and 40 mM chloroacetamide by boiling at 95°C for 10 minutes followed by probe sonication1 second on, 1 second off for 1 minute. Each sample was digested using the SP3 method (1). Carboxylate-functionalized speedbeads (GE Life Sciences) were added to the lysates. Addition of acetonitrile to 80% (v/v) caused the proteins to bind to the beads. The beads were washed twice with 80% (v/v) ethanol and twice with 100% acetonitrile. Proteins were digested with Lys-C/Trypsin (Promega) overnight rotating at 37°C. Speedbeads were collected by centrifugation and then placed on a magnet to remove the digested peptides. The peptides were then desalted using an Oasis HLB cartridge (Waters) according to the manufacturer’s instructions and dried in a speedvac.

Samples were suspended in 3% (v/v) acetonitrile/0.1% (v/v) trifluoroacetic acid and 1 µg tryptic peptides were directly injected onto a C18 1.7 µm, 130 Å, 75 µm X 250 mm M-class column (Waters), using a Thermo Ultimate 3000 RSLCnano UPLC. Peptides were eluted at 300 nL/minute using a gradient from 2% to 20% acetonitrile over 100 min into a Q-Exactive HF-X mass spectrometer (Thermo Scientific). Precursor mass spectra (MS1) were acquired at a resolution of 120,000 from 350 to 1500 m/z with an automatic gain control (AGC) target of 3E6 and a maximum injection time of 45 milliseconds. Precursor peptide ion isolation width for MS2 fragment scans was 1.4 m/z, and the top 12 most intense ions were sequenced. All MS2 spectra were acquired at a resolution of 15,000 with higher energy collision dissociation (HCD) at 30% normalized collision energy. An AGC target of 1E5 and 100 milliseconds maximum injection time was used. Raw files were searched against the Uniprot Varroa destructor database UP000594260 and Apis mellifera database UP000005203 using Maxquant version 2.6.3.0 with cysteine carbamidomethylation as a fixed modification. Methionine oxidation and protein N-terminal acetylation were searched as variable modifications. All peptides and proteins were thresholded at a 1% false discovery rate (FDR).

**S10. Low Temperature Scanning Electron Microscopy.** Low temperature scanning electron microscopy for Figure 1*A-C* was conducted at the USDA-ARS Electron and Confocal Microscopy Unit with techniques outlined by Bolton et al. (2014) and Ramsey et al. (2018). Foundress *Varroa* and eggs were collected from parasitized honey bee brood cells with a fine paintbrush. Each mite was secured to a 15 cm x 30 cm copper plate using ultra smooth, round (12mm diameter), carbon adhesive tabs (Electron Microscopy Sciences, Inc., Hatfield, PA, USA). The specimens were frozen conductively, in a Styrofoam box, by placing the plates on the surface of a pre-cooled (-196 °C) brass bar, the lower half of which was submerged in liquid nitrogen (LN2). After 20-30 seconds, the holders containing the frozen samples were transferred to a Quorum PP2000 cryo-prep chamber (Quorum Technologies, East Sussex, UK) attached to an S-4700 field emission scanning electron microscope (Hitachi High Technologies America, Inc., Dallas, TX, USA). The specimens were etched inside the cryotransfer system to remove any surface contamination (condensed water vapor) by raising the temperature of the stage to -90 °C for 10-15 min. Following etching, the temperature inside the chamber was lowered below -130 °C, and the specimens were coated with a 10nm layer of platinum using a magnetron sputter head equipped with a platinum target. The specimens were transferred to a pre-cooled (-130 °C) cryostage in the SEM for observation. An accelerating voltage of 5kV was used to view the specimens. Images were captured using a 4pi Analysis System (Durham, NC).

Figures

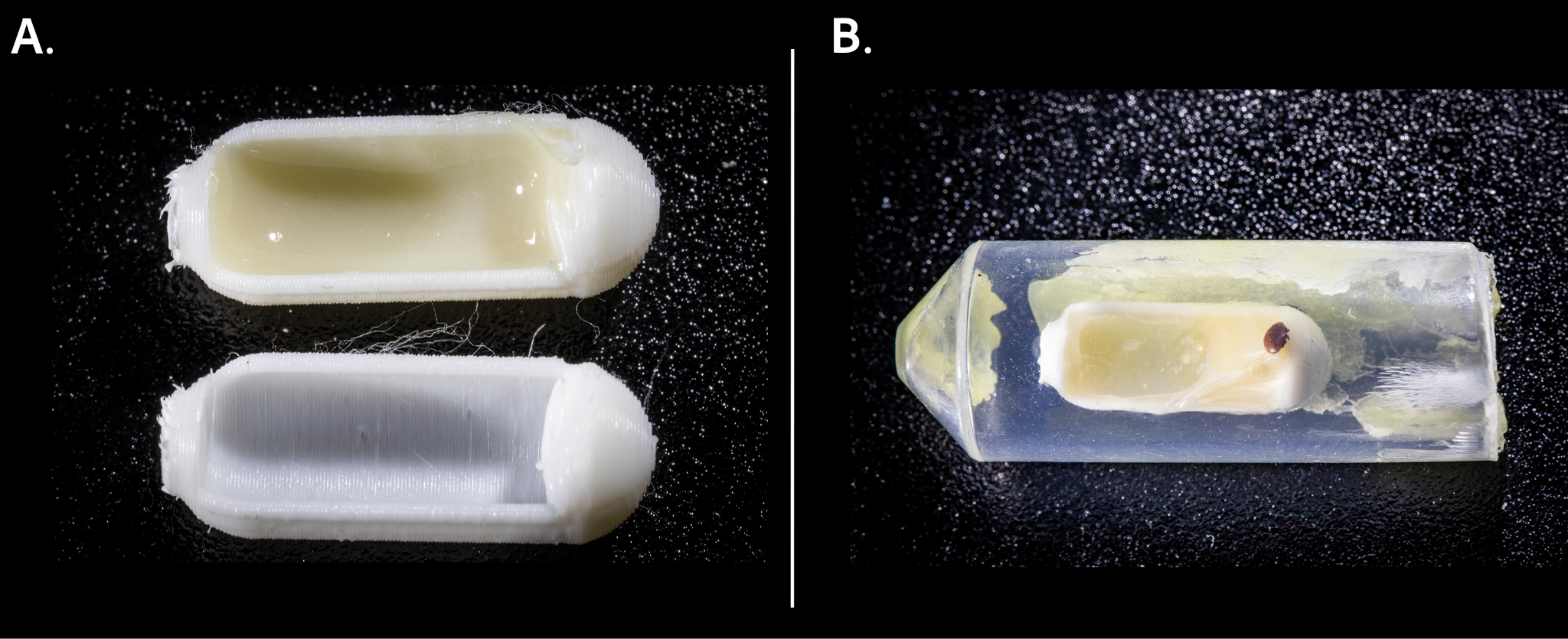

**Figure S1. *Varroa* fluorescent protein feeding trial methods.** (A.) 3D-printed capsules serve as decoy pupae to feed *Varroa* the recombinant Vg-fluorophore complex (Vg-HaloTag-JF549). The inner surface of an empty capsule (lower capsule) and one provisioned with bee tissue (upper capsule) are shown. Provisioned capsules were provided to the control mites containing only a fat body/hemolymph homogenate collected from white-eyed pupae via microdissection while the treatment mites received this mixture with the Vg-fluorophore complex. (B.) The entire *Varroa* feeding apparatus: 3D-printed capsule, food, a layer of 15 µm-thick parafilm, beeswax for adhesion, and a plastic tube to act as the cell. A *Varroa* mite is visible walking on the decoy pupa.

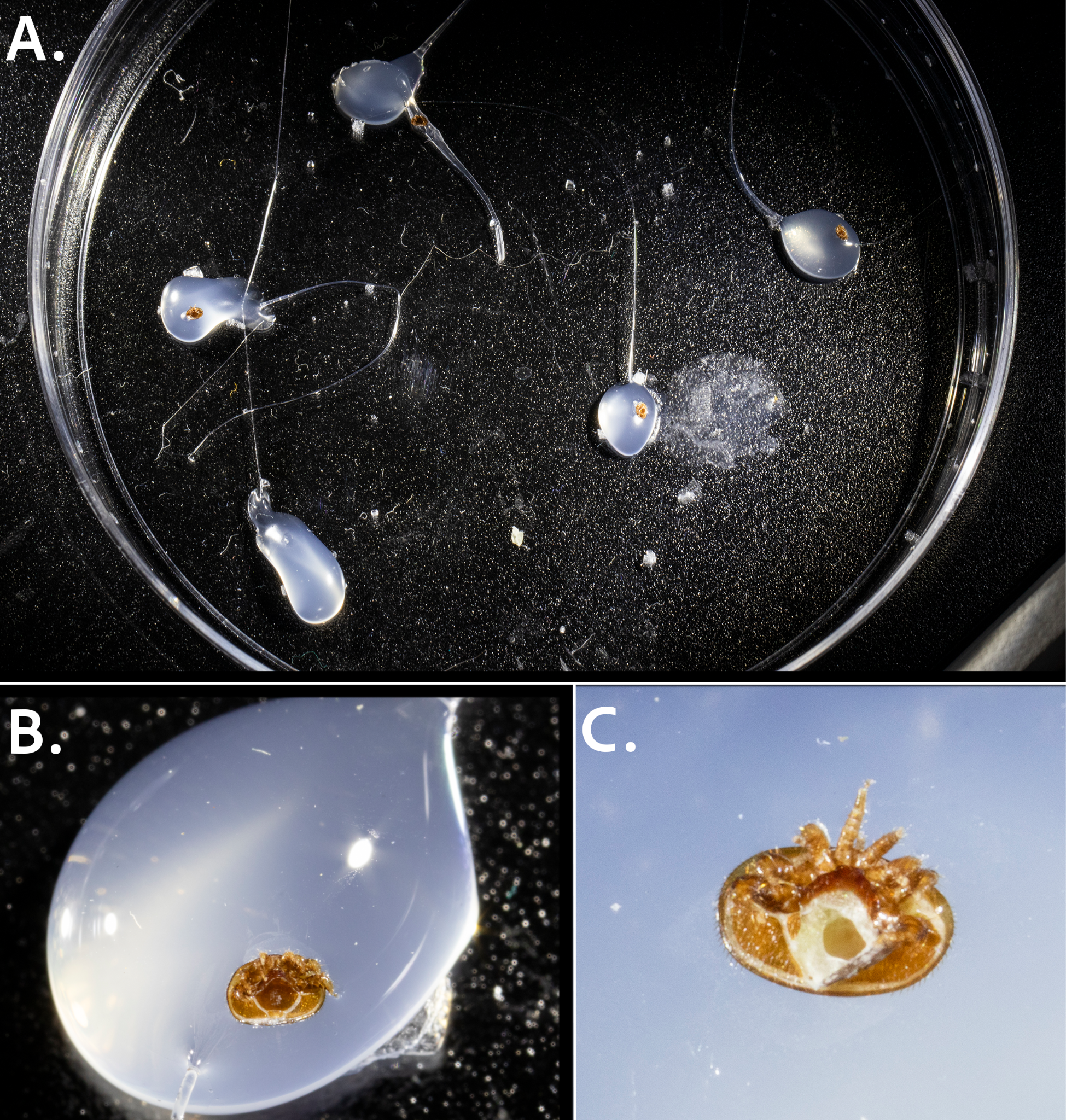

**Figure S2. Mite microdissection methods.** (A.) *Varroa* allowed to feed on fluorophore-laden bee tissue ad libitum were affixed to the lid of a petri dish with hot glue to adhere mites on their dorsal side for microdissection. (B.) Closer image of a *Varroa* mite suspended in glue for microdissection. (C.) A *Varroa* mite after genital plate removal via microdissection. Note that the organs underneath the plate are now visible.

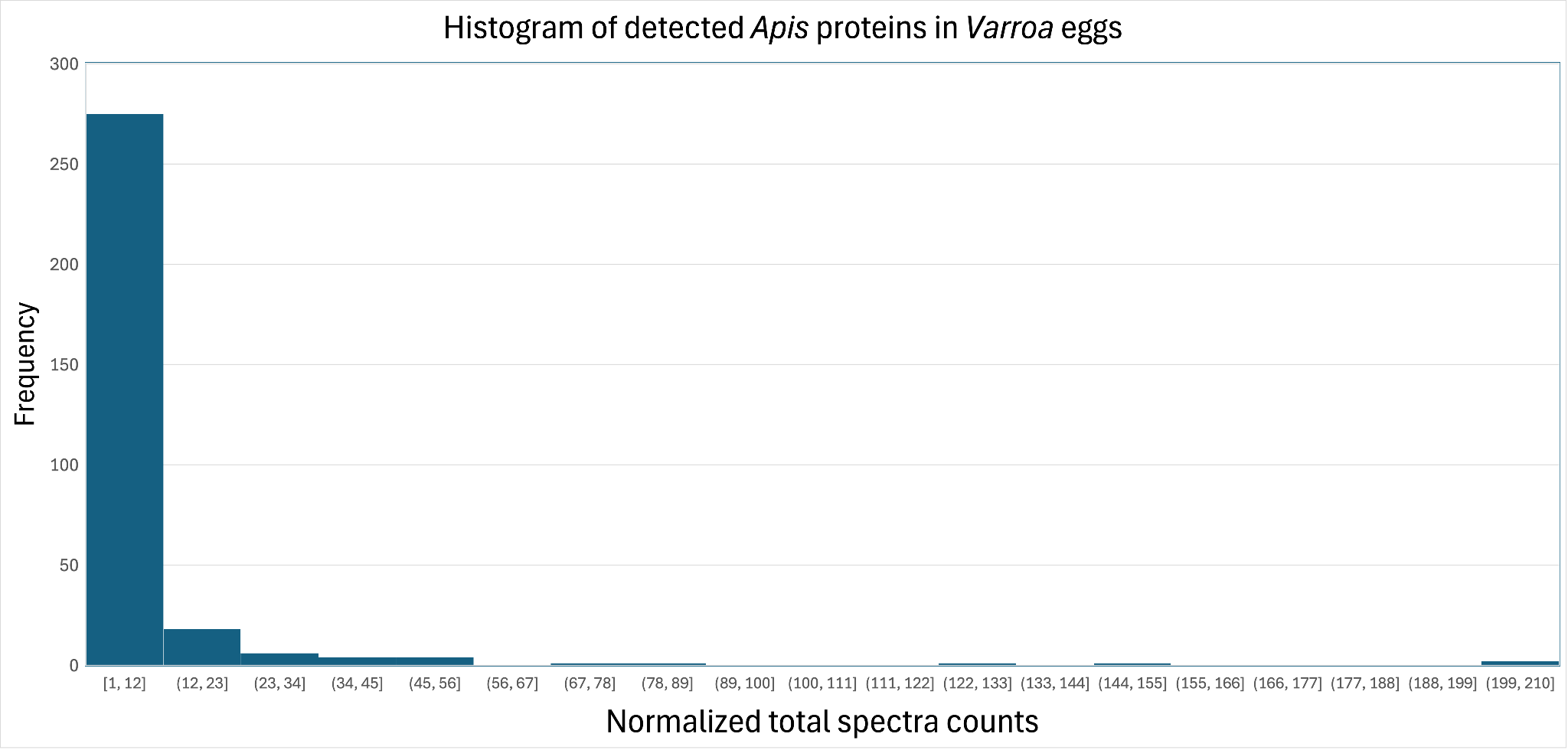
**Figure S3.** Histogram of normalized total spectra counts from *Apis* proteins in *Varroa* eggs. The majority of proteins were detected in low abundance (Normalized total spectrum count < 50).

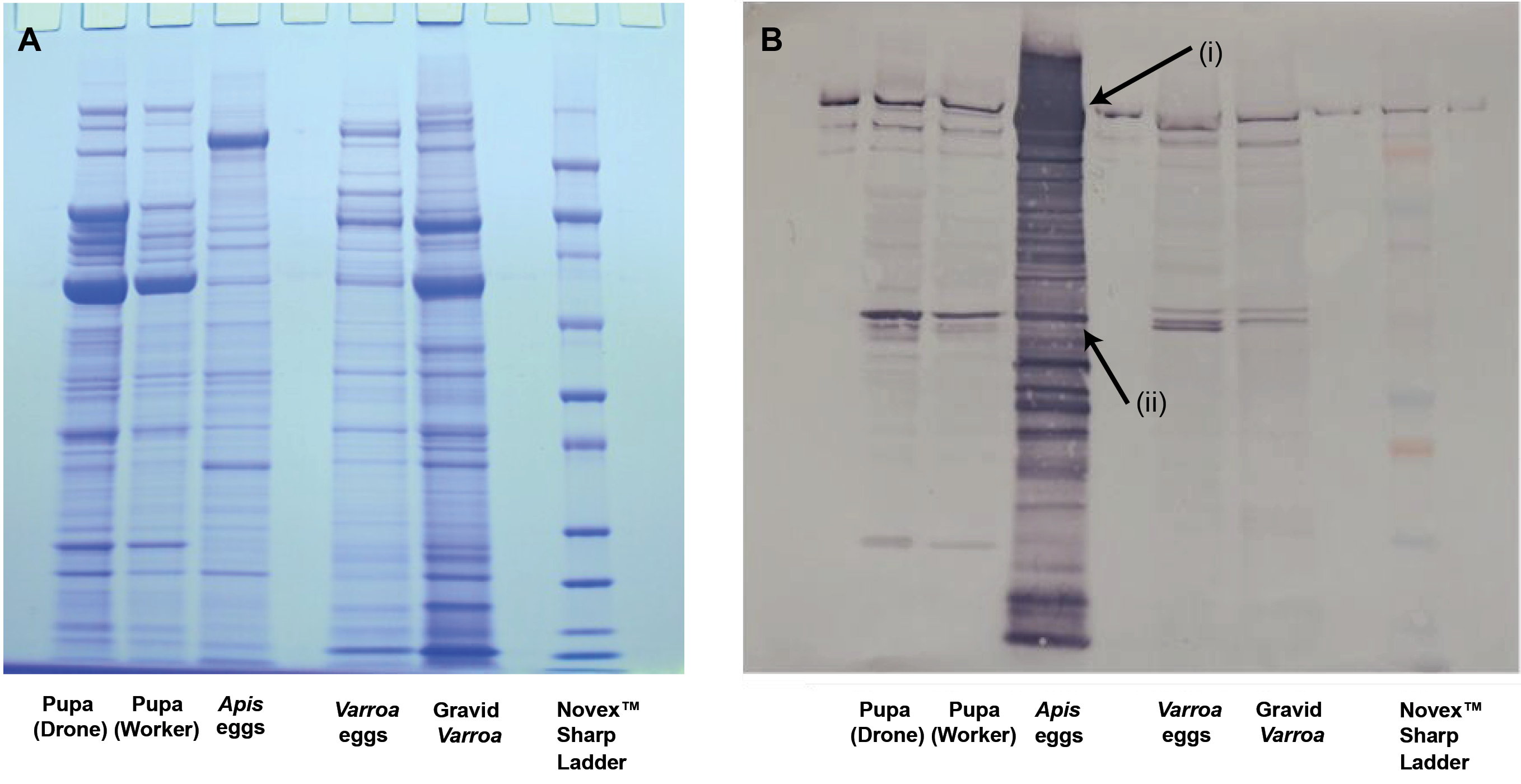

**Figure S4.** SDS-PAGE gel (A.) and Western Blot (B.) showing *Apis mellifera* vitellogenin present in *Apis* and *Varroa* tissue samples. Comparisons shown between proteins from drone and worker honey bee pupae fat body/hemolymph tissue aggregate (*Apis mellifera*), honey bee eggs, and proteins detected in the full body homogenate of parasitic mite *Varroa destructor* (both gravid mites and their eggs). Antibodies were raised to a segment of the protein unique to *Apis Vg* by comparison to the *Varroa* proteome. Arrows indicate vitellogenin bands labeled (i) showing the full-length protein and (ii) a smaller protein which may be a post-translationally modified product (shown to be present in the drone and pupa as well). High-molecular-weight bands visible near the gel wells reflect protein spillover from the highly concentrated *Apis* egg sample. As expected, this signal becomes less distinct farther from the well containing the honey bee eggs. The multi-banded signal in the *Apis* egg lane reflects the high abundance of *Apis* vitellogenin undergoing processing, creating distinct antibody-reactive fragments throughout. As expected, honey bee vitellogenin is detected in honey bee fat body and in especially high abundance in honey bee eggs. This honey bee egg yolk precursor was also detected in *Varroa* and *Varroa* eggs. Note that Varroa eggs display a distinctive band not present in the gravid female mites suggesting further post-translational processing or digestion within the egg.

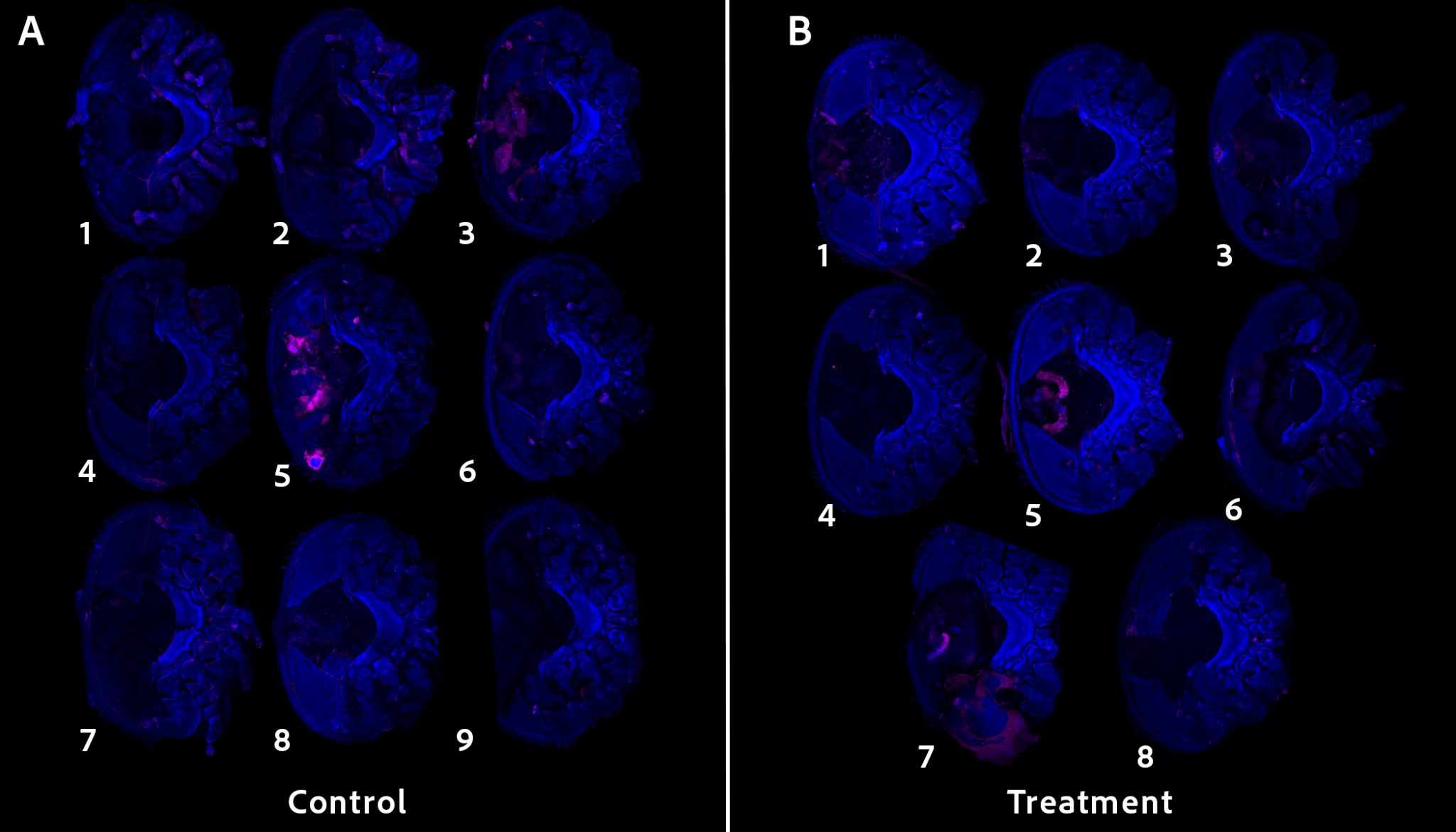

**Figure S5. Full cohort of *Varroa* used in a Vg-complex feeding trial.** Confocal fluorescence microscopy of all control (N = 9) and treatment (N = 8) specimens. (A) Control mites provided homogenized bee tissue without fluorophore ad libitum. Some specimens displayed non-localized background autofluorescence, but lacked the discrete, organ-specific morphology observed in the treatment specimens. (B) Treatment mites provided homogenized bee tissue supplemented with Vg-HaloTag-JF549 to consume ad libitum. Localized fluorescence was observed in the lyrate organ (B, mites 5 and 7) and the lower digestive system (B, mite 5). We removed the genital plate in all specimens prior to imaging. Most mites in the Vg-HaloTag treatment group showed no fluorescence that could be attributed to the fluorophore, likely because they did not feed on our in vitro rearing system and thus did not ingest the fluorophore.

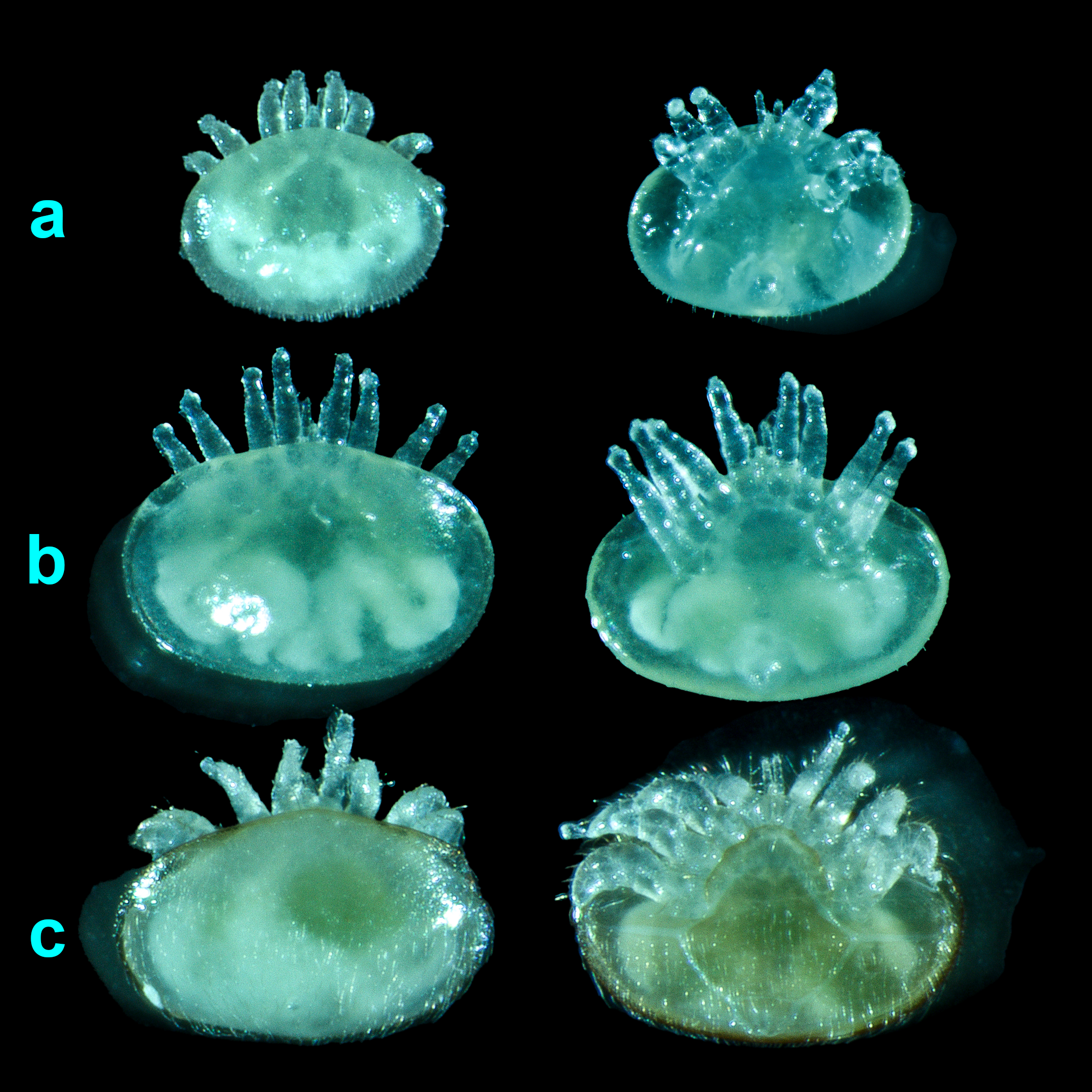

**Figure S6.** **Life stages of *Varroa destructor* collected for LFQ analysis of proteins depleted during metamorphosis.** Images display the dorsal (left) and ventral (right) orientations of developing mites harvested from capped honey bee brood cells. From youngest to oldest, (a) depicts a deutonymph, displaying functional appendages, an unpigmented and pliable exoskeleton, and developing an ellipsoid-shape; (b) depicts a calyptostase nymph, identified by immobility and splayed, turgid appendages; (c) depicts a teneral adult, showing a freshly molted female with fully formed plates and hirsute integument, but still lacking full adult cuticle pigment and sclerotization.

Tables

**Table S1.** Primer sequences, melting temperatures, and amplicon sizes for the three fragments of *Apis* vitellogenin from total cellular RNA.

| **Primer** | **Primer Sequence (5’-3’)** | **T_m_** | **Amplicon Size** |
| --- | --- | --- | --- |
| Fragment 1 F | ATGTTGCTACTTCTAACGCTTTTACT | 65°C | 1746 bp |
| Fragment 1 R | GTTCACTTGGGCATTGTGGA |  |  |
| Fragment 2 F | AAGAGGAGCATTCACAATAATTATCCAG | 65°C | 1666 bp |
| Fragment 2 R | ACGTGGAGAATATCACAGGATCTTT |  |  |
| Fragment 3 F | CGTTCGACAGCAAGGTGATG | 67*C | 1898 bp |
| Fragment 3 R | AGCCTTGCAAACGAAAGGAAC |  |  |

**Table S2**. Energy content of components during *Varroa’s* reproductive stage, acquired through bomb calorimetry and used to create the static energy budget.

| **Sample** | **Sample mass (g)** | **Wet unit mass (mg)** | **Energy per unit (J)** |
| --- | --- | --- | --- |
| *Varroa* eggs | 0.0129 | 0.043 | 0.379 |
| Reproductive *Varroa* | 0.1 | 0.395 | 2.503 |
| Dispersing *Varroa* | 0.068 | 0.300 | 2.606 |
| Emerging *Varroa* | 0.036 | 0.360 | 1.555 |
| Just-capped *Varroa* | 0.0047 | 0.490 | 2.555 |
| *Varroa* feces (30 hours) | 0.06 | 0.133 | 1.828 |
| *Apis* fat body (30 hours) | 0.2 | 1.100 | 5.226 |

**Table S3. Independent micro-computed tomography (microCT) volume measurements for *Varroa destructor* adults and eggs.** Measurements were taken for non-gravid, adult foundress mites (N = 10) and eggs (N = 10) collected from capped honey bee brood cells. Based on the ratio of the means (0.05297 mm^3^ egg/0.3017 mm^3^ adult), a single egg constitutes approximately 18% of the adult’s body volume.

| **Adult Mites (N = 10)** | **Volume (mm^3^)** | **Eggs (N = 10)** | **Volume (mm^3^)** |
| --- | --- | --- | --- |
| 1 | 0.2893 | 1 | 0.04261 |
| 2 | 0.3146 | 2 | 0.05572 |
| 3 | 0.3308 | 3 | 0.05591 |
| 4 | 0.3486 | 4 | 0.04391 |
| 5 | 0.2120 | 5 | 0.04592 |
| 6 | 0.2044 | 6 | 0.03937 |
| 7 | 0.3439 | 7 | 0.05690 |
| 8 | 0.3467 | 8 | 0.03951 |
| 9 | 0.3706 | 9 | 0.06992 |
| 10 | 0.2561 | 10 | 0.07994 |
| **Mean Adult Volume** | 0.3017 | **Mean Egg Volume** | 0.05297 |
| **SD** | ± 0.05917 | **SD** | ± 0.01354 |
| **Range** | 0.2044 - 0.3706 | **Range** | 0.03937 - 0.07994 |

**Table S4.** Statistical analysis of hexamerin 110 log_2_ fold-change across developmental stages. Log_2_ fold-changes of hexamerin 110 (HEX110) were compared to the mean log_2_ fold-change of background proteins across three life stage transitions. Background protein fold-change variance was calculated as ±1 standard deviation (S.D.) from the mean.

|  | **LFC** | **P - adj.** |
| --- | --- | --- |
| **Deutonymph – Teneral Adult** | | |
| Hexamerin 70b | -2.811 | 0.042 |
| Hexamerin 70c | -3.654 | 0.010 |
| Hexamerin 110 | -4.192 | 0.023 |
| Other Proteins  (mean ± S.D.) | -2.152 ± 1.424 |  |
| **Deutonymph:** **Calyptostatic** | | |
| Hexamerin 70b | -0.479 | 0.508 |
| Hexamerin 70c | -1.772 | 0.040 |
| Hexamerin 110 | -3.161 | 0.117 |
| Other Proteins (mean ± S.D.) | -1.435 ± 1.132 |  |
| **Calyptostase – Teneral Adult** | | |
| Hexamerin 70b | -2.299 | 0.134 |
| Hexamerin 70c | -1.873 | 0.176 |
| Hexamerin 110 | -1.066 | 0.543 |
| Other Proteins (mean ± S.D.) | -0.740 ± 1.120 |  |

**Table S5. Results of permutation tests comparing metamorphic hexamerins (110, 70b, and 70c) to other host proteins in immature mites.** Permutation tests were performed with 100,000 permutations to generate the null distribution.

| **Transition** | **Mean difference**  **(hex110, 70b, 70c - other proteins)** | **P-value** |
| --- | --- | --- |
| **Deutonymph – Teneral** | -1.399654 | 0.04253957 |
| **Deutonymph – Calyptostase** | -0.3738757 | 0.2680573 |
| **Calyptostase – Teneral** | **-**1.000341 | 0.0701993 |

**Table S6. Comparison between the reproductive mass between *Triatoma phyllosoma* and *Varroa destructor*.** Values for *T. phyllosoma* are from Collier et al. (1977). We assume that an egg’s wet mass is 70% water (Maino et al. 2016). Reproductive investment is defined as the amount of eggs produced at one time. We use the average clutch size for *T. phyllosoma* (13.3 eggs) to represent a single reproductive investment of *Triatoma*, and that each egg is 3.36 mg (wet mass). *Varroa* only produces one egg at a time, thus its reproductive investment is 1 (Ifantidis, 1997). By mass alone, *Varroa creates* a larger reproductive investment than *T. phyllosoma*.

| Species | Adult wet mass | Body mass increase for reproduction | Reproductive investment (R.I.), wet mass | R.I. /Adult mass |
| --- | --- | --- | --- | --- |
| *Varroa destructor* | 0.300 mg  (dispersing) | 31.6% | 0.043 mg | 14.3% |
| *Triatoma phyllosoma* | 373.24 mg | 71.9% | 44.77 mg (13.3 eggs, 3.36 mg per egg) | 11.9% |

**Table S7. Peptides used to sensitize rabbit hosts for antibody collection.** Honey bee and *Varroa* Vg showed 17.2% sequence similarity, providing an abundance of potential target sequences.

| **Name** | **Peptide** | **Position** | **Length** |
| --- | --- | --- | --- |
| OBS-603:620 | DNSLYDEYIPFLERELRK | 603 | 18 |
| OBS-743:760 | LMKLKSPEWKDLAKKARS | 743 | 18 |
| OBS-768:782 | HEYDYELSRGYIDEK | 768 | 15 |
| OBS-1264:1277 | KPKMDFNVDIRYGK | 1264 | 14 |
| OBS-1283:1300 | ERIDMNGKLRQSPRLKEL | 1283 | 18 |
| OBS-1738:1755 | KLKKRIEKGANPDLSQKP | 1738 | 18 |

**Table S8. Local BLASTp alignment between *Apis* vitellogenin antibody sequence and endogenous *Varroa destructor (Vd)* vitellogenins.** High E values confirm the unlikelihood of antibody cross reactivity between species’ vitellogenins.

| Antibody Sequence | Target Protein | NCBI Accession No. | Max Score | Total Score | Query Cover Representative Sequence | E Value | Per. Ident |
| --- | --- | --- | --- | --- | --- | --- | --- |
| KLKKRIEKGANPDLSQKP | *Vd* Vg1 | AFN88463.1 | 9.5 | 18.7 | 22% | 41 | 100.00% |
|  | *Vd* Vg2 | AFN88464.1 | 10.0 | 29.1 | 44% | 29 | 100.00% |

Dataset S1 (separate file). Full list of all *Apis-* and *Varroa-* derived proteins identified through high-performance liquid chromatography-tandem mass spectrometry. Samples consisted of excised gel bands from mite eggs, reproductive mites, dispersing (non-reproductive) mites, adult bee hemolymph, adult bee fat body, and immature bee pupal hemolymph/fat body aggregate. All samples were normalized in Scaffold (Proteome Software) using the standard spectrum count normalization algorithm, in which spectral counts for each sample were scaled by the ratio of the mean total spectral count across all samples to the spectral count of the individual sample. Normalized spectral counts from each gel band from a given tissue (e.g., mite eggs) were summed to provide the total spectral counts per sample.

Dataset S2 (separate file). Full List of all 313 *Apis-* derived proteins in Varroa eggs. Proteins were identified through high-performance liquid chromatography-tandem mass spectrometry. Samples consisted of excised gel bands from mite eggs, reproductive mites, dispersing (non-reproductive) mites, adult bee hemolymph, adult bee fat body, and immature bee pupal hemolymph/fat body aggregate. All samples were normalized in Scaffold (Proteome Software) using the standard spectrum count normalization algorithm, in which spectral counts for each sample were scaled by the ratio of the mean total spectral count across all samples to the spectral count of the individual sample. Normalized spectral counts from each gel band from a given tissue (e.g., mite eggs) were summed to provide the total spectral counts per sample. *Apis*- derived proteins which were not found in *Varroa* eggs are excluded, as are *Varroa*- derived proteins.
